# Dissecting the molecular origins of opalescence and phase separation in mAb formulations and their relation to aggregation

**DOI:** 10.1101/2025.08.29.672998

**Authors:** Inna Brakti, Samuel Lenton, Marco Polimeni, Hannes Ausserwöger, Rob Scrutton, Anette Henriksen, Nikolai Lorenzen, Martin Nors Pedersen, Jesper Søndergaard Marino, Fátima Herranz-Trillo, Ann E. Terry, Tuomas P. J. Knowles, Minna Groenning, Vito Foderà

**Affiliations:** Department of Pharmacy, University of Copenhagen, Copenhagen, Denmark 2100; Biophysical Analysis, CMC Analytical Support, Novo Nordisk A/S, Måløv, Denmark 2760; Yusuf Hamied Department of Chemistry, Centre for Misfolding Diseases, University of Cambridge, Cambridge, United Kingdom CB2 1EW; Novo Nordisk A/S, Therapeutics Discovery, Novo Nordisk Park, Måløv 2760, Denmark; MAX IV Laboratory, Lund University, Fotongatan 2, 224 84 Lund, Sweden

**Keywords:** antibody development, patchy interactions, liquid-liquid phase separation, opalescence, aggregation

## Abstract

Liquid-liquid phase separation (LLPS) and high opalescence are two self-association phenomena commonly encountered in monoclonal antibody (mAb) formulations. Because of their impact on colloidal stability, they are commonly avoided, due to a suspected link with aggregation and reduced product shelf-life. However, the molecular underpinnings and interrelation between these phenomena remain unclear, complicating predictions of their occurrence. By combining light and X-ray scattering techniques with microscopy and advanced microfluidic setups, we here report the delicate phase behavior of a model mAb, named mAb1. This is characterized by rapid clustering and LLPS in a narrow NaCl range, above which it transitions into an opalescent state devoid of micron-sized assemblies, yet retaining a similar interaction fingerprint. Using Monte Carlo simulations, we report that the macroscopic solution state of mAb1 is controlled by a positive patch, whose degree of charge screening determines whether LLPS or opalescence will take place. Specifically, neutralization of this patch via counterion interactions diminishes intermolecular repulsion and favors the concerted action of weaker dipole-dipole/hydrophobic interactions, amounting to the creation of a new solution phase, via LLPS. Further NaCl addition distributes ions more uniformly across the surface, attenuating these attractive interactions, leading to the dismantling of droplets while preserving solution opalescence. Finally, we show that LLPS and opalescence are decoupled from stirring-induced aggregation, challenging an unequivocal relationship between these phenomena.

**Significance Statement:** Tailoring formulations to maximize the stability of therapeutic antibodies is crucial for their development. This is complicated by their tendency for self-association at high concentrations, where increased opalescence and phase separation, that are thought to precede irreversible aggregation, are routinely observed. Here, we studied the molecular underpinnings of mAb opalescence versus liquid-liquid phase separation. We report the mechanisms determining the two phenomena and provide a foundation for their prediction, which may guide the rational development of mAb formulations. We further show that LLPS and opalescence can be decoupled from stress-induced aggregation. We hypothesize that excluding mAbs from the bulk solvent via LLPS may even be harnessed to enhance drug product stability.

## Introduction

Since the approval in 1986 by the US Food and Drug Administration of the first monoclonal antibody (mAb) as a novel drug modality, their commercial success has been continuously expanding (1–4). Therapeutic mAbs are usually administered parenterally by intravenous infusion or subcutaneous (SC) injection. For chronic disease management, the latter approach is favored, as it improves patient adherence, by providing a less intrusive delivery method and the ease of self-administration (5–8). However, due to their large size and often limited potency, mAbs formulations can exceed 100 mg/mL to reach relevant doses in the small injection volumes imposed by the SC route (≈2mL) (8, 9). In these concentrated regimes, the probability of collisions between mAb molecules increases and can result in the formation of higher-order protein assemblies. This can have several consequences on solution properties such as increased viscosity (10), opalescence (11) liquid-liquid phase separation (LLPS) (12) and/or the formation of soluble/insoluble aggregates (13). Opalescence refers to increased light scattering as a result of reversible attractive interactions (14) and is associated with large concentration fluctuations in the vicinity of a phase transition boundary without macroscopic assembly as detected by light microscopy (11, 15). This leads to a bluish solution appearance, akin to an opal stone (11). LLPS, on the other hand, is a phenomenon in which attractive interactions between macromolecules outbalance solvent interactions and physically separate from the bulk solution into a discrete phase with liquid-like properties (16). This results in macromolecules concentrating into droplets, forming the “dense” phase, surrounded by a macromolecule-depleted “dilute” phase (16). However, as the properties of the droplet interior can be inhomogeneous, the more inclusive term “phase-separation” is preferably employed, unless the liquid-like nature of the droplets is confirmed (17). Both opalescence and LLPS are reversible and dependent on solution conditions. Despite the solution factors inducing these phenomena being well-known, much less is known about the molecular mechanisms driving opalescence versus LLPS. This is complicated by the fact that pure opalescence and LLPS can be visually hard to distinguish. The latter involves the formation of a new phase, often witnessed as micron-sized droplets which are detectable via microscopy, while high opalescence can be observed without any large-scale assembly. In a drug development context, opalescence above a defined threshold is often avoided as it is commonly flagged as a precursor for LLPS (6). The latter is a concern for two reasons: (1) the delicate formation and dissociation of droplets, based on changes in temperature, excipients, protein concentration and incubation time complicates the development of proper control strategies during a product’s shelf-life and (2) it has been linked to aggregation (11, 18). LLPS-related aggregation is mostly reported for protein systems with some level of intrinsic disorder (19–23). This has often been observed as a liquid-to-solid transition within “aged” droplets (24). Moreover, the distinct environment provided by the droplet interface has been shown to serve as nucleation point for fibril formation for several intrinsically disordered proteins (17, 22, 25). The importance of this inherent property of many proteins was highlighted for the amyotrophic lateral sclerosis-related protein SOD1, whose exposure of disordered loops in the absence of zinc led to phase separation and subsequent fibril formation (23). Interestingly, while droplet interfaces have been demonstrated to spearhead aggregation, recent reports also point toward a shielding role of the droplet interior, emphasizing the double-edged nature of this phenomenon (26, 27). Concerning large, folded mAbs, Pantuso et al. hypothesized that the LLPS-induced partial unfolding of a mAb promoted crystal growth along the droplet surface (28). On the other hand, Bramham et al. observed that the structural integrity of another mAb was conserved upon LLPS formation and no changes in the aggregation profile were detected after storage at refrigerated conditions (4°C) after 28 days (29).

In this study we integrated light- and X-ray scattering and microscopy, to discriminate between mAb opalescence and LLPS. By combining these approaches with controlled microfluidic mixing, we accessed a wide design space to carefully monitor the progression of interactions underlying LLPS, as well as their inhibition. Using coarse-grained Monte Carlo (MC) simulations, we rationalized the microscopic differences between LLPS and opalescence in terms of molecular interactions leading to the two phenomena. Finally, we linked this fundamental study of the macroscopic manifestations of mAb self-assembly to effects on the physical stability under stress conditions to support future formulation studies of mAbs.

## Results

### mAb1 opalescence and liquid-liquid phase separation are modulated by ionic strength

We started out by investigating the phase behavior of 45 mg/mL mAb1 samples in 10 mM histidine pH 6.5, with increasing NaCl concentration (0-150 mM) at room temperature (RT). Inspection of the solutions revealed an increase in the solution opalescence (Figure 1A) and right-angle light scattering (RALS) intensity (Figure 1B) at low salt, followed by a less steep decrease at higher NaCl concentrations, until returning to a clear solution at 150 mM NaCl. To determine the origin of this high opalescence (which we will simply refer to as opalescence from now on), we next investigated selected mAb solutions by brightfield microscopy, which are highlighted in colors in Figure 1B. Images of the 0 mM and 150 mM NaCl samples with no solution opalescence were clear under the microscope (Figure 1C). However, samples at 10 and 22.5 mM NaCl were characterized by distinct solution morphologies, despite comparably high scattering intensities and similar visual appearance. At 10 mM NaCl, spherical droplets, with liquid-like properties were observed (see Figures 1C and S1), while increasing the NaCl concentration to 22.5 mM abolished this phenomenon. The presence of both opalescence as a single phenomenon (14) or associated with LLPS in a narrow NaCl range complicates the determination of the LLPS regime by e.g. turbidity measurements (16). We therefore combined simultaneous microfluidic mixing and fluorescence detection of LLPS events for the high-throughput mapping of the mAb1 phase diagram at low NaCl concentrations and over a broad mAb1 concentration range (30). Briefly, this method relies on the precise encapsulation of predefined combinations of mAb1, buffer and NaCl solutions into microenvironments surrounded by an oil phase, for which the presence of LLPS is individually assessed via fluorescence detection. As depicted in Figure 1D (and Figure S2-S3), adding as little as 5 mM NaCl induced LLPS of mAb1 at concentrations above an approximate saturation concentration of 30 mg/mL. Remarkably, above 10 mM NaCl, the system exited the de-mixed regime and underwent reentrant phase transition (31, 32) into a homogeneous state, characterized by high light scattering intensities and absence of droplets.

**Figure 1.**
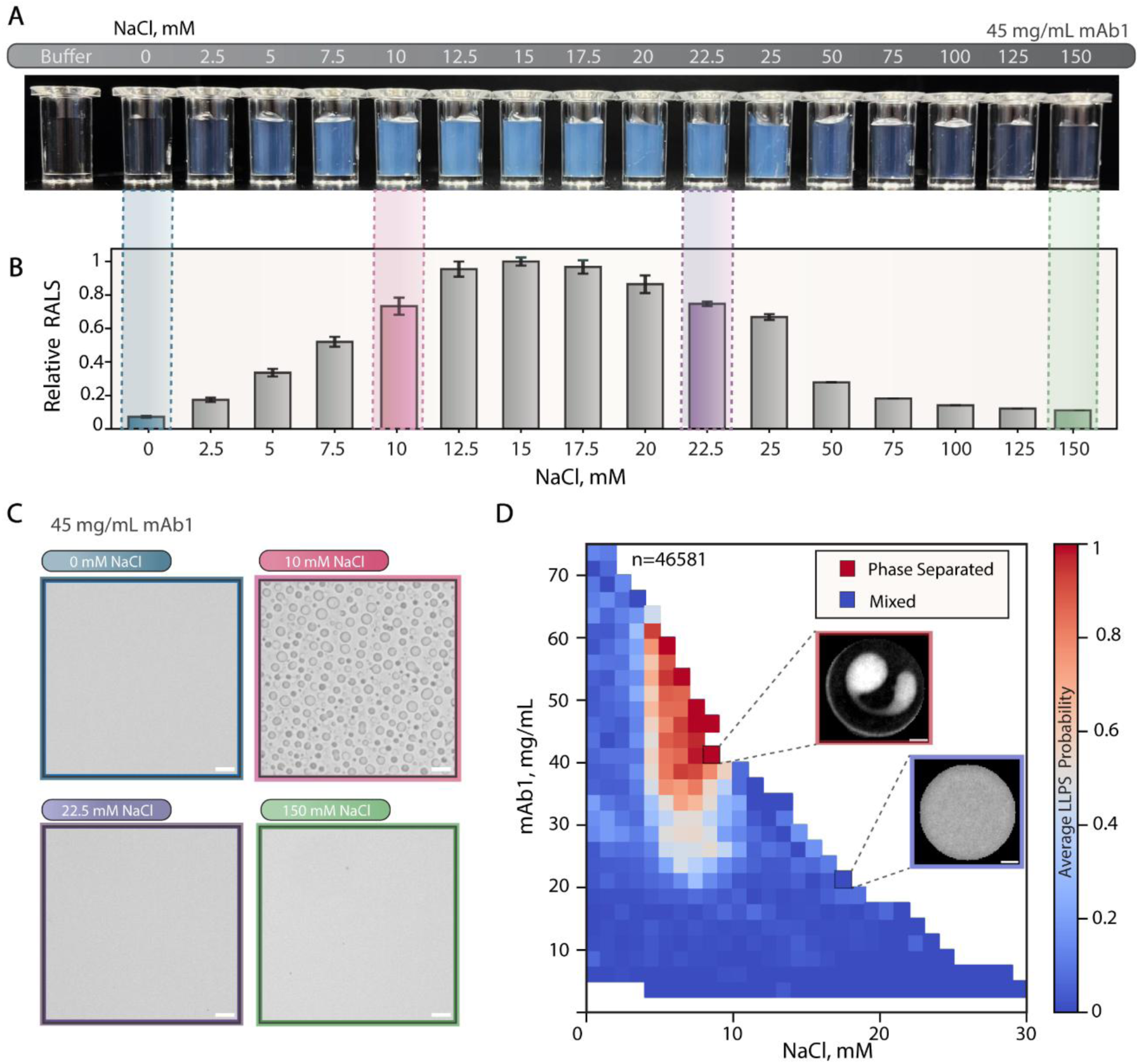
Phase behavior of mAb1. (A) Visual appearance and (B) normalized right-angle scattering intensities at 635 nm of 45 mg/mL mAb1 solutions as a function of NaCl, where selected conditions are highlighted in colors. The experiments were independently performed n=2 times and errors represent standard deviations. (C) Brightfield microscopy images of selected conditions (0, 10, 22.5 and 150 mM NaCl, 20 µm scalebar). (D) Phase diagram of mAb1 versus NaCl concentration obtained by measuring 46581 independent data points with a combinatorial microfluidic platform (70) (LLPS probability = 1: red, 0.5: white, 0: blue). Representative microscopy images of phase-separated (red)and mixed (no LLPS, blue) are included as insets.

### Opalescence and LLPS have similar interaction fingerprints

To differentiate between opalescence and LLPS, we studied the protein-protein interactions (PPIs) governing these phenomena, by measuring the small- and wide-angle X-ray scattering (SAXS and WAXS) profiles of our selected formulations at 45 mg/mL. Drastic differences in the scattering intensity I(**q**) were observed in the low **q** region of the scattering curves (larger distances), while the curves overlapped in the high **q** region (Figure 2A). Both the LLPS (10 mM NaCl) and opalescent (22.5 mM NaCl) samples were characterized by highly attractive interactions, as inferred from the upturn of the scattering intensity I(**q**) at low **q**, compared to the form factor (FF) obtained in the absence of PPIs (black in Figure 2A, S4). This feature was, however, more prominent for the LLPS condition. A common strategy to minimize such attractive interactions in mAb formulations relies on the addition of L-arginine HCl (hereafter referred to as arginine) (9, 33). Indeed, adding 150 mM arginine to a formulation in which mAb1 would have otherwise undergone LLPS (i.e. 45 mg/mL mAb1 + 10 mM NaCl in 10 mM histidine, pH 6.5), no LLPS was observed (Figure S5A). This was also the case for the opalescent sample (Figure S5B). When comparing the effect of adding arginine versus 150 mM NaCl, which similarly decreased solution light scattering, the former resulted in a SAXS/WAXS scattering curve overlapping with the FF, while the latter presented a deviation therefrom, indicating residual attractive forces in solution.

**Figure 2:**
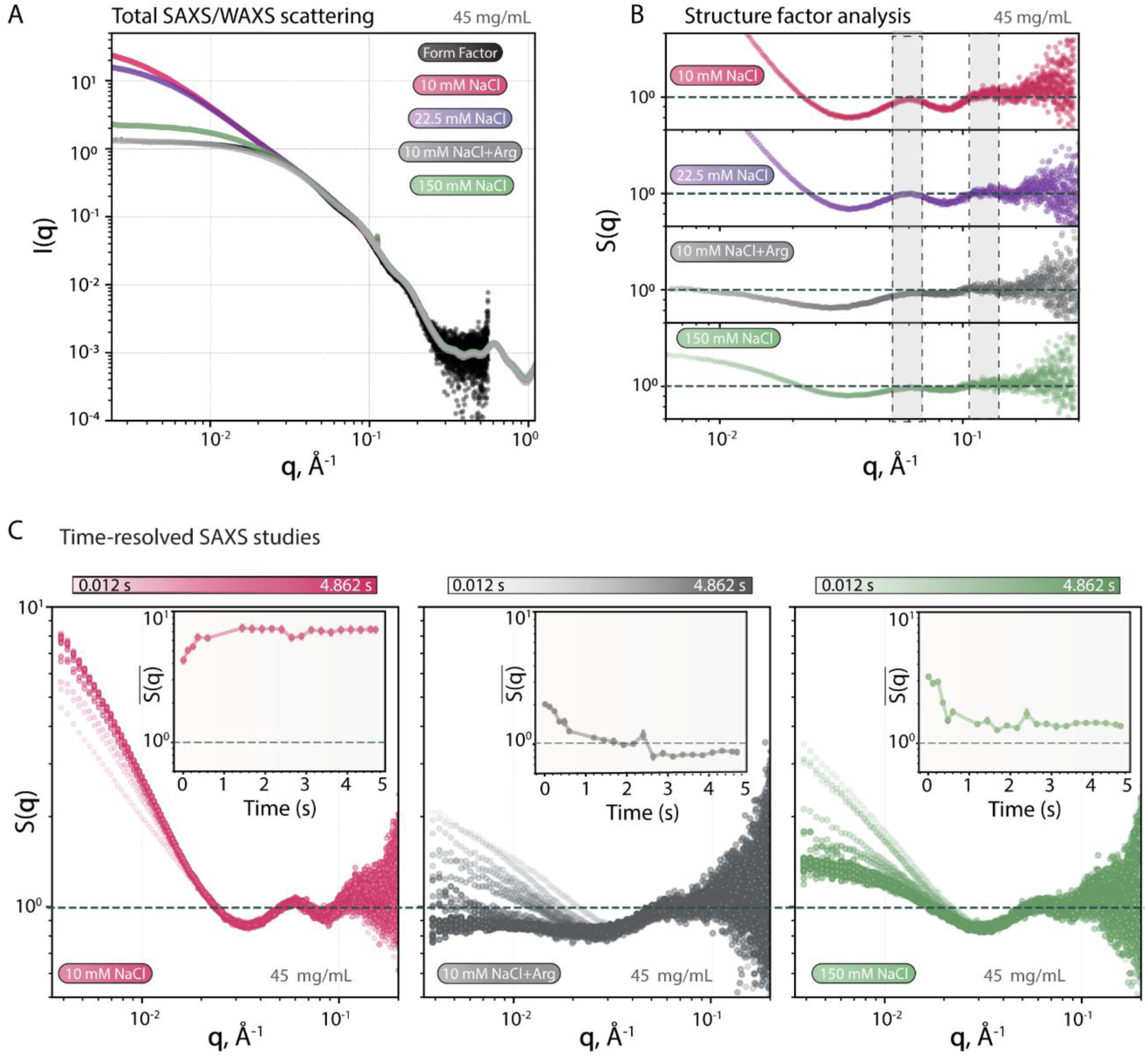
Characterization of interactions underlying mAb1 LLPS and opalescence with SAXS. (A) SAXS/WAXS intensities of 45 mg/mL mAb1 + 10 (pink), 22.5 (purple) or 150 mM NaCl (green) and 10 mM NaCl + 150 L-arginine HCl (grey, called 10 mM NaCl+Arg here) as well as the form factor obtained from SEC-SAXS (black) (B) Structure factors measured for the same samples (C) Time-resolved SAXS data (21 datapoints spanning 0.012-4.862s) of 45 mg/mL+ 10 mM NaCl (pink), 10 mM NaCl+arg (grey) and 150 mM NaCl (green). Averages of the first three S(**q**) values for each curve as a function of time from mixing are included as insets for a clearer overview of the evolution of PPIs.

We next extracted structure factor contributions, S(**q**), from the total scattering intensity, to emphasize the nature of PPIs at our selected conditions (Figure 2B). For both the LLPS (pink) and opalescent (purple) conditions, this analysis revealed a large positive slope in S(**q**) at low **q**. Furthermore, positional correlation peaks were observed at **q**=0.06 Å^-1^ and 0.1-0.12 Å^-1^, corresponding to real space distances of 104 and 52 Å, i.e. the diameter and radius of a mAb (34). Such an S(**q**) profile is reconcilable in terms of the formation of reversible mAb clusters in solution (5, 35). Interestingly, the same peaks were also visible for the condition containing 150 mM arginine (termed 10 mM NaCl+Arg here, grey) and 150 mM NaCl (green) but with decreased peak definition, probably due to the stronger screening of electrostatic interactions at higher ionic strength (IS) (36).

To study the progression of PPIs involved in cluster formation and LLPS, we added a temporal dimension to our SAXS experiment, by coupling a microfluidic device to the beamline as described in (37) and illustrated in Figure S6. Under LLPS-inducing conditions for mAb1, the above-mentioned fingerprint for cluster formation was already visible 12 ms after the mixing point and increased slightly until stabilizing after ≈1.5s (Figure 2C, pink). Addition of arginine or 150 mM NaCl conditions gradually decreased these interactions, albeit more slowly in the latter case, despite the lower molecular weight (Mw) of NaCl. In all cases, the scattering curves of the final timepoints in the time-resolved experiment overlapped with the static SAXS data, showing a high level of agreement between the two setups (Figure S7). Altogether, this suggests that both LLPS and opalescence have similar underlying molecular fingerprints, characterized by attractive interactions and cluster formation, which can be inhibited using similar strategies.

### mAb1 phase behavior is dictated by electrostatic patch-mediated interactions

To dissect the molecular features driving LLPS at 10 mM, opalescence without phase separation at 22.5 mM and a transparent solution at 150 mM NaCl, we employed Monte Carlo (MC) simulations. We first matched the experimental scattering curves, I(**q**), using a 174-bead coarse-grained model of mAb1. We tuned our model to reproduce the strength of the short-range hydrophobic attractions, (*ε*_*ij*_ in equation 2, Materials and Methods), using the I(**q**) curve corresponding to 150 mM NaCl, where we expected hydrophobicity to be the most prominent, due to the electrostatic screening (green in Figure 3A). This could be achieved at *ε*_*ij*_=0.3185 kJ/mol. However, the simulated I(**q**) curves for the 22.5 mM and 10 mM NaCl at pH 6.5 showed a less attractive behavior compared to the SAXS data (Figure 3A). The structures used for the simulations at pH 6.5 are shown in Figure 3B. Their 1 kBT/e electrostatic surface potentials at 10, 22.5 or 150 mM NaCl show a highly anisotropic charge distribution, and we estimated a high positive net charge of the mAb (Figure S8). Considering the latter and the sole bulk screening effect of NaCl, a high degree of repulsion should be indeed expected at low salt, which is gradually screened at higher salt concentrations. This is also captured by our two-body simulations in terms of calculated mAb reduced second osmotic virial coefficient, B2*, from the potential of the mean force (PMF in Figure S9), and the electrostatic multipole expansion (Figure S10). The decreasing B2* values obtained at pH 6.5 at increasing NaCl concentrations in Figure 3C originate from a progressively screened repulsion, mirroring the trend observed for the simulated I(**q**) (Figure 3A).

**Figure 3:**
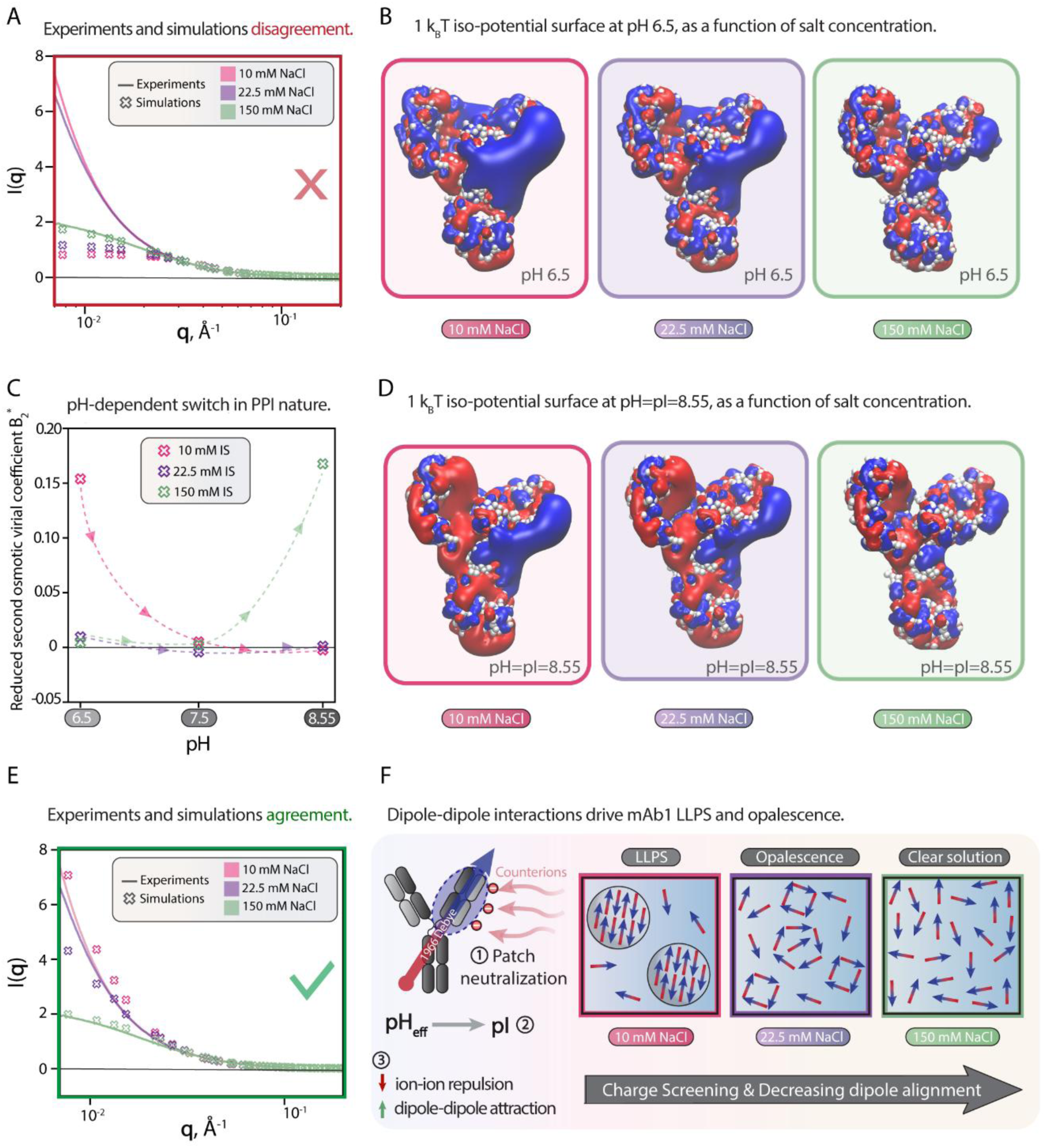
**Monte Carlo simulation of mAb1**. (A) SAXS intensities, of a 45 mg/mL mAb solution at pH 6.5 obtained from experiments (full lines), and simulations (small *x* symbols) of 100 mAbs coarse-grained with the 174-bead model for different IS (B) Reduced osmotic second virial coefficient, B2*, of mAb1 obtained from two-mAb simulations at different pH and IS. 1 *kBT/e* electrostatic surface potential of mAb1 at (C) pH 6.5 and (D) pH 8.55 for three different IS: 10, 22.5 and 150 mM (left to right) obtained with adaptive Poisson-Boltzmann solver (APBS). Blue indicates a positive electrostatic surface, while red indicates a negative electrostatic surface. The dipole moment of mAb1 was calculated to be 1966.64 Debye at 10 mM NaCl, pH 6.5. (E) Same as (A) but at pH=8.55=pI. (F) Summarizing illustration of molecular mechanism leading to different phenomena based on charge screening effects after the pH shift. At 10 mM NaCl (pH 6.5) interactions are repulsive due to the high net charge of mAb1. As a lot of this excess positive charge is accumulated in a patch in one of its Fab domains, a small addition of NaCl screens this patch to an extent where the overall charge state of mAb1 is neutralized, akin to what it would be close to its pI. Therefore, solution counterions can modulate the local pH experienced by proteins. This suppression of ion-ion repulsion allows mAb1 dipoles to align in a configuration favorable for LLPS. Further charge screening suppresses LLPS at 22.5 mM NaCl but preserves some level of dipole alignment. While the properties of the resulting clusters do not allow them to phase-separate, they nonetheless result in a high level of solution opalescence. Moving even further away from the phase boundary by adding 150 mM NaCl results in an even lower probability of mAb1 dipole aligning, which is witnessed as a clear solution characterized by more repulsive interactions.

To rationalize the inverse trend observed experimentally, a key aspect needs to be considered. Prominent positive surface patches (blue in Figure 3B) are known to be involved in mAb self-assembly (38–41). In general, electrostatic patches can interact with salt ions and effectively alter the solution pH experienced locally by moving it towards the isoelectric point (pI) (42). Following this rationale, we evaluated the B2* at pH values approaching the pI of mAb1. At pH 7.5, B2* nearly overlaps at all salt conditions, indicating a similar interaction profile (Figure 3C). At the pI (≈8.55), however, the order inverts: the 10 mM salt curve (pink) exhibits the strongest attraction, followed by 22.5 mM (purple), while the highest salt concentration (green) is the least attractive, recapturing the experimental trends. Looking at the multipole decomposition analysis (Figures S10-S12), this trend is explained by a gradual decrease in the repulsive ion-ion interactions from pH 6.5 to 8.55, until vanishing completely and leaving only attractive dipole–dipole contributions. The presence of the mAb dipole moment at pH 8.55 is evident from the electrostatic isopotential surface (Figure 3D), showing a higher symmetry between the positive (blue) and negative (red) potentials that are gradually screened with increasing salt. We then performed many-mAbs simulations using the 174-bead model and calculated the I(**q**) curves at pH 8.55. As shown in Figure 3E, our simulations at pH 8.55 successfully recapitulate the experimental trend, strongly supporting a mechanistic scenario in which salt-induced saturation of the mAb1 electrostatic patch drives a pH shift toward the pI. This shift plays a pivotal role in enhancing the contribution of attractive hydrophobic and dipole–dipole interactions, ultimately promoting LLPS at 10 mM NaCl. Upon further salt addition to 22.5 mM NaCl, the strength of dipole–dipole interactions diminishes due to dipole screening (43) and the dense phase is disrupted, but substantial mAb–mAb attraction is maintained, which manifests macroscopically as opalescence without phase separation. At 150 mM NaCl, extensive dipole screening results in reduced inter-mAb attractions.

### LLPS/opalescence and stress-induced aggregation of mAb1 are decoupled

Finally, we investigated the link between LLPS and opalescence in mAb formulations and propensity for stress-induced aggregation. We therefore subjected the same formulations (45 mg/mL mAb1, 10, 22.5, 150 mM NaCl or 10 mM NaCl+Arg) to vigorous stirring (16 hours at 1000 rpm, RT). This resulted in a dramatic turbidity increase in all formulations (Figures 4A and 4B). Interestingly, arginine did not protect against contact stirring despite its reported role as a suppressor of aggregation (44). The visibly aggregated samples were centrifuged for 2 hours at 21.000 *g*, resulting in pellets comparable in size across the different formulations. Strikingly, the 10 and 22.5 mM NaCl samples stayed opalescent after stirring. Moreover, the hard centrifugation revealed the presence of a transparent top dilute phase as previously observed for mAb network formation (45). We next verified by brightfield microscopy that aggregates were responsible for the observed pellet (Figure 4C). Interestingly, droplets with preserved liquid-like properties were observed in the soluble fraction of the stirred 10 mM NaCl sample, demonstrating that LLPS persists after harsh stirring or can reform rapidly thereafter (Figure 4C, bottom). Finally, we calculated the monomer recovery in the different formulations. To avoid density gradients from different phases in the LLPS/opalescence samples, we added 150 mM arginine directly to the samples before and after stirring, when measuring their concentrations (Figure 4D). Within the error, no differences in monomer recovery between the samples were detected, suggesting that the stirring-induced aggregation observed is independent of IS (Figure 4E). Aggregation was observed even after the formulations were supplied with 0.15 mg/mL polysorbate 80 (PS80) indicating that the observed phenomena do not solely arise from the air-water interphase but also from the bulk solution (Figure S13).

**Figure 4.**
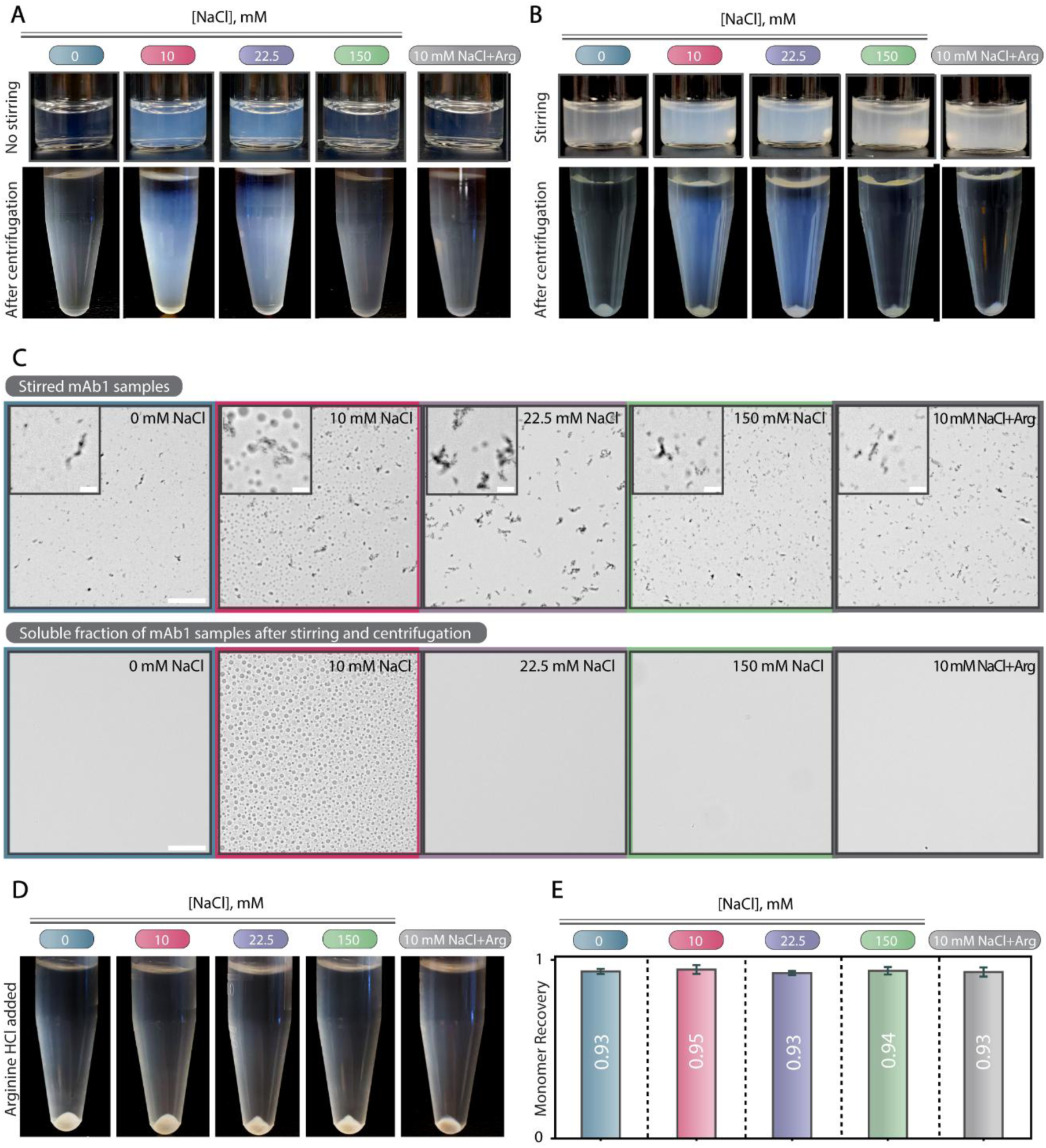
Physical (in)stability of mAb1 samples as a result of stirring stress. Visual appearance of 45 mg/mL mAb1 solutions at pH 6.5 containing 0, 10, 22.5, 150 mM NaCl or 10 mM NaCl+Arg before (A) and after (B) magnetic stirring (16 hours) at 1000 rpm and subsequent centrifugation. (C) Brightfield microscopy of the stirred samples and the supernatant after centrifugation (50 µm scalebar). (D) Representative images of the stirred and centrifuged samples with added L-arginine HCl. Samples were prepared at a larger volume for the images for increased clarity. (E) Monomer recovery of the different conditions measured on two independent samples. The numbers on the individual bars represent the calculated monomer recovery, where 1 and 0 stand for complete and no monomer recovery, respectively. Errors represent standard deviations from duplicate measurements.

**Figure 5:**
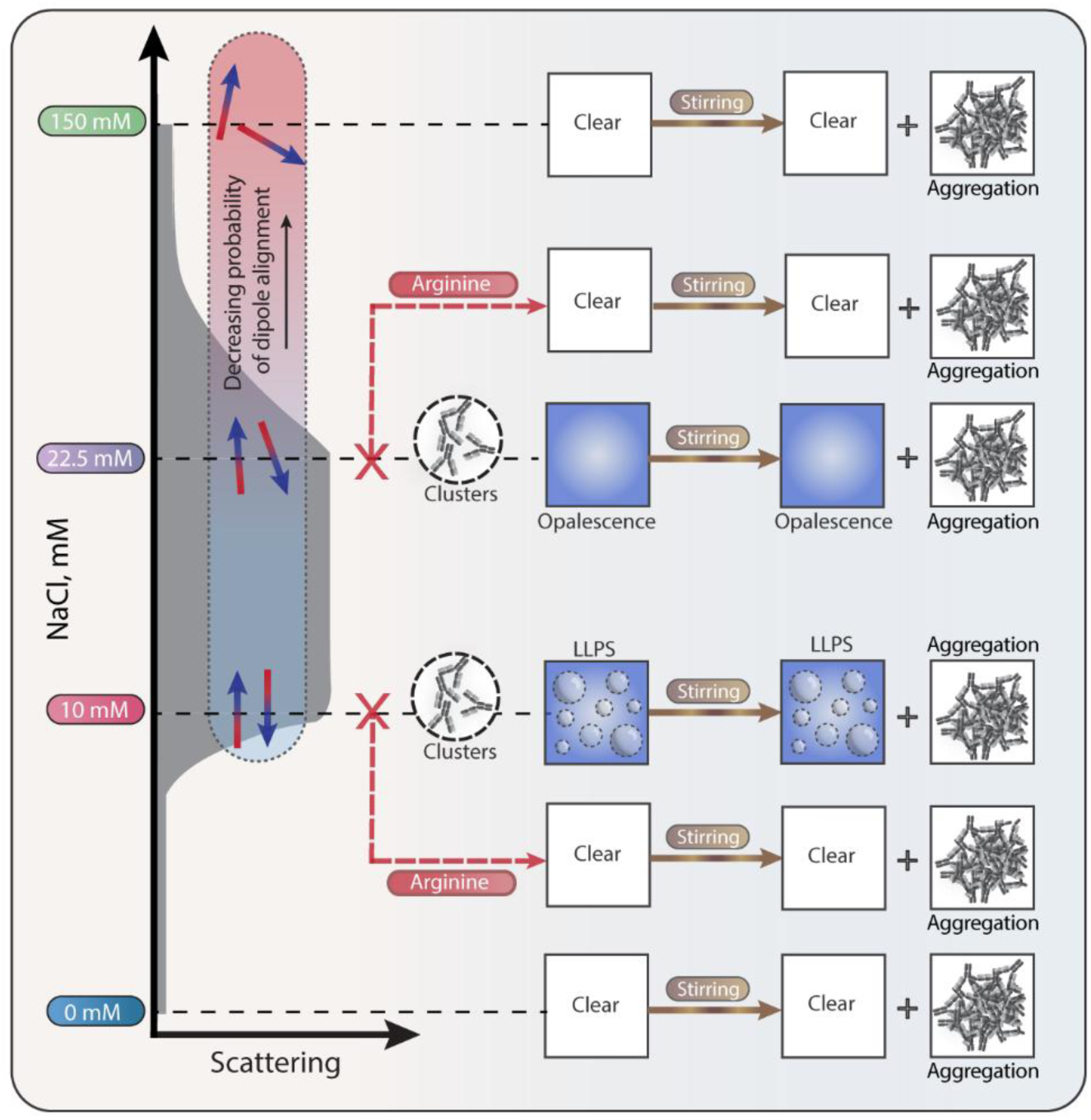
Summary of mAb self-assembly behavior at 45 mg/mL, as a function of NaCl concentration and applied stirring stress. mAb1 LLPS and opalescence occur in a narrow NaCl range can both give rise to similar light scattering intensities, despite having different underlying assembly levels. Both phenomena are characterized by attractive interactions, leading to cluster formation, which can be suppressed by arginine. The level of charge screening in solution dictates the probability of mAb1 dipoles to align favorably and results in large scale association, which decreases with NaCl concentration. Solid aggregation is decoupled from LLPS and opalescence. LLPS and opalescence are unaffected by stirring induced stress.

## Discussion

Here, we studied the model antibody mAb1, who undergoes LLPS above a concentration of ∼30 mg/mL, in a narrow range of low NaCl concentrations in 10 mM histidine buffer at pH 6.5 (Figure 1). Above this NaCl range, the mAb1 solution returns to a one-phase state, characterized by a highly opalescent, droplet-free phase, in the vicinity of the upper critical NaCl concentration for LLPS.

In our SAXS/WAXS studies (Figure 2), we noted a shared interaction fingerprint between LLPS and opalescence, characterized by highly attractive interactions, leading to cluster formation. These were shown to occur within milliseconds upon mixing components, leading to mAb1 LLPS, and could be efficiently mitigated by arginine, resulting in a clear solution (Figure 2 and S5). We observed that addition of 150 mM arginine led to a higher degree of inhibition of these interactions compared to adding 150 mM NaCl. Due to comparable ionic strengths, this indicates that the effect of arginine is not purely ionic, as noted by Tilegenova et al. (46). Notably, the scattering patterns at high **q** values (small distances) overlapped with the form factor both for the LLPS and opalescent conditions, suggesting that these phenomena are not accompanied by large structural rearrangements. Importantly, the macroscopic differences between these two phenomena imply that their molecular underpinnings differ. The sharp response in phase behavior to a slight change in IS as well as the abrogation of LLPS and opalescence further pointed towards key interactions being mainly confined to oppositely charged surface patches (5, 39).

To clarify this, we performed simulations to reproduce the trends in our experimental X-ray scattering data and obtain a model describing the interactions driving LLPS versus opalescence. The simulations only agreed with the experimental phase behavior when shifting the solution pH towards the pI of mAb1 (see Figure 3A and Figure 3E). This indicates a mechanism in which the highly positive net charge of mAb1 is progressively neutralized while adding salt, leading to an effective shift of the mAb pH towards the pI. Our electrostatic isopotential surface analysis (Figure 3B) has shown the presence of a strong positive electrostatic patch ranging from the Fab to Fc domains in mAb1. We hypothesize that in the presence of a small amount of salt (10 mM of NaCl) the electrostatic path gets neutralized, and this causes the pH to shift towards the pI. At the pI, the suppression of the strong ion-ion repulsions allows short-ranged attractive dipole–dipole and hydrophobic interactions to become more prominent, shifting the balance towards a net attraction of mAbs in solution. In this configuration dipole-dipole interactions are strong, as dipoles can align to maximize their interaction energy. This process is at the basis of the formation of stable clusters, leading to droplet formation and LLPS (Figure 3F) (43). The mechanism of the patch neutralization and the consequent shift towards the mAb pI is in line with the work of Pineda et al. (42), who demonstrate that patchy distributions of ionizable groups on protein surfaces significantly affects pH in protein solutions, linking local charge inhomogeneities to deviations between bulk buffer pH and the effective pH experienced by proteins. Similarly, Ausserwöger et al. observed that FUS and PGL3 condensates minimized electrostatic repulsion by buffering towards the pI of the molecules, via the exclusion of buffer molecules (47).

When a higher amount of salt is added to the solution (22.5 mM of NaCl) the electrostatic patch is neutralized, and the excess ions distribute uniformly over the mAb surface. This causes a partial screening of the dipoles and a consequent misalignment. This translates into weaker dipole-dipole attractions and a consequent incapacity of the system to phase-separate yet preserving some alignment among dipoles and giving rise to the high opalescence (Figure 3F). Clusters responsible for this phenomenon are likely smaller or less stable, compared to those witnessed during LLPS (48). This also explains the small decrease observed in the low-**q** region of the scattering intensity at 22.5 mM NaCl (Figure 2A). At 150 mM NaCl, electrostatic screening becomes much stronger and dipole–dipole interactions are negligible, leaving hydrophobic interactions as main attractive force between mAbs. This makes the solution appear clear as both opalescence and LLPS are disrupted.

Finally, we questioned whether LLPS and opalescence of mAb solutions could lead to increased aggregation (Figure 4). We chose to evaluate this in a forced degradation setup where formulations were subjected to stirring stress, as this has been shown to rapidly induce the formation of large amounts of aggregates for mAbs (49–52). In this setup, stirring of all mAb1 formulations produced solid pellets of comparable sizes, suggesting that its effect is independent of IS in the timeframe studied (Figure 4B). Importantly, LLPS was conserved at 10 mM NaCl, coexisting with solid aggregates (Figure 4C). This was also the case for samples containing PS80, suggesting that the bulk solution also contributes to the overall aggregation (Figure S13). The fact that hard centrifugation couldn’t completely sediment LLPS droplets suggests that their volume fraction is small and that there could exist an equilibrium between them and clusters in solution, as shown for other systems (17, 24, 53). Finally, the observation that LLPS and opalescence were unaffected by stirring and that the same amount of mAb material was converted into aggregates in all formulation points towards the presence of two decoupled mechanisms. One in which mAb1 self-association is dependent on IS, leading to charge modulation of the mAb1 surface in such a way that weak attractive interactions can amount to LLPS or opalescence, depending on the level of charge screening. The other stress-induced mechanism is independent of IS (and hence LLPS/opalescence), resulting in the formation of solid aggregates. Hence, mAb1 LLPS and opalescence are not inherently associated with an increased proclivity for aggregation in our accelerated setup. Whether the effects observed under conditions of harsh stirring translate into long-term stability during, for example, storage remains an important question for future investigation. Nevertheless, the observation that LLPS of mAb1 remains unaffected by aggregation following harsh stirring stress suggests that this phenomenon is stable within mAb formulations. This stability could potentially be harnessed to protect mAbs from other manufacturing stresses or to enable formulations at even higher concentrations. More quantitative, experimental studies, combined with simulations (25) may help in uncovering possible liquid-to-solid transitions over longer timescales under these conditions.

Taken together, our data highlight the diverse phase behavior of mAb1, characterized by LLPS at low salt, followed by reentrant phase transition to a highly opalescent state, both driven by attractive interactions and cluster formation. To our knowledge, the present study represents one of the very few studies showing that monovalent ions can induce reentrant phase transitions in mAb formulations. We propose that localized interactions between surface patches and solvent ions modulate the effective pH experienced by mAb1 and steer it towards a more charge neutral state, where dipole-dipole and hydrophobic interactions can dominate and induce LLPS or opalescence, depending on the level of dipole screening by NaCl. Finally, LLPS and opalescence have traditionally been viewed as precursors of aggregation in mAb formulations. Here, we report a not so straightforward relation between these phenomena, which should be taken into consideration during the rational design of mAb formulations.

## Materials and Methods

### Model antibody

The model mAb used in this work, mAb1, was supplied by Novo Nordisk A/S in a proprietary formulation before being stored at -20°C as aliquots.

### Buffer and stock preparation

A buffer containing 10 mM L-histidine was prepared freshly using reagent grade L-histidine (ReagentPlus®, ≥99% TLC, H8000, Sigma Aldrich) in Milli-Q water and the pH was adjusted to 6.5. Sodium chloride (1.37017.5000, Merck), L-arginine HCl (EMPROVE® EXPERT, Ph. Eur., ChP, JP, USP, 1.01587, Sigma Aldrich) and polysorbate 80 stocks (1.37171, high purity, EMPROVE® EXPERT, Ph. Eur., ChP, JP, NF, Sigma Aldrich) were prepared in buffer and filtered through a 0.1 um Whatman Anotop filter (GE Healthcare, Cat no: 6809-2012).(53)

### Buffer exchange by dialysis

Two rounds of dialysis were carried out using Slide-A-Lyzer™ cassettes (A52976, Merck, 20 kDa Mw cutoff) in buffer for two hours at RT (22°C) and one round overnight (16 hours) at 4°C using fresh buffer with a volume 500 times larger than the sample volume at each step.

### Sample concentration determination

Protein concentrations were determined as independent duplicates by measuring the UV-Vis absorbance at 280 nm on a NanoDrop™ One C spectrophotometer. Serial 1:1 dilutions of the mAb stock solution in buffer were prepared from a factor of 2 to 64. The absorbance of each dilution was measured using a theoretical molar extinction coefficient at 280 nm and the Mw before being multiplied by the dilution factor.

### Right-angle static light scattering

Static light scattering experiments were performed at 90° with a 635 nm semiconductor laser (Labbot spectrophotometer, ProbationLabs, Sweden). Samples were transferred to a high precision quartz glass, 10x4 mm light pathlength cuvette (104F-10-40, Hellma Analytics, Germany). Measurements were performed at RT, on independent duplicate samples. These were filtered through a 0.1 um Whatman Anotop filter (GE Healthcare, Cat no: 6809-2012). As soon as the scattering signal stabilized, ten consecutive measurements were performed before being averaged. The data was normalized to the highest scattering intensity.

### Brightfield microscopy

Brightfield microscopic images were acquired at RT with a Zeiss Axioscope 5 equipped with an Axiocam 305 color camera, through an LD Plan-Neofluar 20x/0.4 objective and processed using ImageJ software (54).

### Combinatorial microfluidic mixing (PhaseScan)

The phase space of mAb1 as a function of NaCl was mapped out at room temperature using combinatorial microfluidic mixing setup, according to published procedures (30, 47). Pre-defined combinations of Alexa647 N-hydroxy succinimidyl ester labeled mAb1, 10 mM histidine buffer (pH 6.5) and a 40 mM NaCl stock containing Alexa488 dye were encapsulated in HFE-7500 oil (FluoroChem) supplied with 1.2% (w/V) fluorosurfactant, (RAN Biotechnologies) by flowing the latter into the microfluidic device at a constant flowrate. A total of 46581 droplets containing pre-defined combinations of aqueous components were evaluated for the presence of LLPS (Figures S2 and S3) using an inverted fluorescence microscope (OpenFrame, Cairn Research, Faversham, UK) equipped with a camera (Kinetix, Photometrics, Tucson, AZ, USA, 20x objective). The fluorescence intensities in the separate channels were used to back calculate the mAb1 and NaCl concentrations for each droplet. A grid-map visualization was achieved by binning the individual data points as described in (47).

### Synchrotron size-exclusion chromatography (SEC)-SAXS for form factor determination

SEC–SAXS experiments were performed at the P12 EMBL BioSAXS beamline, at PETRA III, EMBL Hamburg, operated at a fixed energy (10 keV, λ = 0.124 nm). The form factor was determined at room temperature by SEC-SAXS using a Superdex S200 column coupled to an Agilent1260 Infinity Bio-Inert high performance liquid chromatography (HPLC) system with an in-built temperature-controlled auto-sampler. A total volume of 70 µL mAb1 solution at a concentration of 10 mg/mL in 10 mM histidine pH 6.5+150 mM NaCl was injected into the column. The same buffer was used as mobile phase, and the flowrate was set to 0.65 mL/min. SAXS data was collected for the whole column run. The resulting frames were processed using CHROMIXS from the ATSAS package (55). See Figure S4 for the SEC-SAXS chromatogram and resulting form factor data. By fitting the Guinier region (56) of the obtained form factor, we determined the radius of gyration of mAb1 to be 50 Å, as expected for a standard antibody (34).

### SAXS/WAXS

SAXS and WAXS data of mAb1 at 45 mg/mL in 10 mM histidine pH 6.5 containing 10, 22.5 and 150 mM NaCl or 10 mM NaCl+ 150 mM L-arginine HCl at room temperature was collected at the P12 (PETRA III, DESY synchrotron, Hamburg) and CoSAXS beamlines (MaxIV laboratory, Lund University), following standard procedures. Frames of the measured total scattering intensity I(**q**) as a function of the momentum transfer **q** were averaged and buffer subtracted. The curves were then absolute scaled in ATSAS (57), using water and empty capillary measurements, performed under the same conditions. The structure factor S(**q**) was obtained by dividing the scaled buffer-subtracted scattering curve with the form factor.

### Time-resolved solution SAXS

Time-resolved solution SAXS was performed at the CoSAXS beamline at the MAX IV Laboratory, using a microfluidic sample delivery system (35). Briefly, solution stocks (or buffer) are added to the inlets of a microfluidic-chip, with three inlets (cyclo-olefin copolymer COC/TOPAS®, 300x400 µm cross-section, Microfluidic ChipShop). These a mixed at a mixing point to reach the condition of interest, after which SAXS is recorded at different locations in the channel.

Two inlets contained 60 mg/mL freshly dialyzed mAb stock, while the last inlet in the middle position was supplied with either buffer or excipient stock solutions (NaCl and/or L-Arginine HCl) stocks. At the mixing point, the mAb is diluted by a factor of 2/3 and the excipients, by a factor of 1/3. SAXS data was collected at two of the inlets prior to mixing, the mixing point and 21 positions from the mixing point at a total flowrate of 60 µL/min as the chip was translated along **z**. The time resolution of the experiment was determined by a combination of the flowrate and the distance from the mixing point. SAXS data was recorded with an incident X-ray beam of wavelength λ = 0.99 Å, with a sample-to-detector (Eiger2 X 4m) distance of 3.5 m yielding a q-range of 0.004 < **q**< 0.3 Å^−1^. The background of each inlet was measured prior to replacing the contents of the inlet 1 and 3 with mAb solution. 40 frames were recorded at each point for a total of 50 ms and averaged. The scattering curves were absolute scaled using water and empty capillary measurements performed under the same conditions. All processing was performed using the Primus package in the ATSAS data analysis software (57) and S(**q**) was calculated as described in the previous section.

### Structure preparation for Monte Carlo simulations

A structural model of mAb1 was generated by replacing the Fab domains in a public IgG4 crystal structure (PDB:5DK3) with those of mAb1. Using the Visual Molecular Dynamics visualization program (58), the mAb sequence was extrapolated. The conformational energy was minimized using OpenMM (version 8.3) (59), as follows. Missing hydrogen atoms were added to reflect the protonation state at pH 6.5. The system was then parametrized using the Amber14 ff14SB (60) force field for proteins, combined with the TIP3Pwater model (61). The Particle Mesh Ewald method (62)was employed, to deal with long-range electrostatics. Using the Langevin integrator (63) at 300 K, iterative energy minimizations were executed, using OpenMM’s L-BFGS optimizer (64), to reduce potential energy by removing steric clashes and relaxing strained geometries. Starting from the high positive potential energy we reached a minimum plateau after performing 1000 steps of maximum 100 iterations.

### Coarse-graining and charge calculation

From the atomistic energy-minimized structure of mAb1, a coarse-grained (CG) model at the amino acid (aa) level was generated, by replacing each aa with a single bead, positioned at its center of mass. To each bead was assigned an effective diameter, *σ*_*i*_ according to the equation:

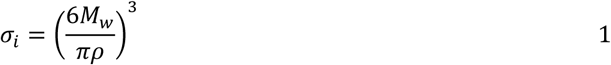

where *ρ* = 1 g/mol is an average aa density (65) and *M*_*w*_, the mass of the single aa. To accurately assign aa charges under varying solution conditions, constant pH Monte Carlo (MC) simulations were performed, using Faunus (66, 67). To each bead was assigned an intrinsic pKa value (68) reflecting the acid-base equilibrium constant of the isolated side chain under ideal solvent-exposed conditions (66). These values underpin the protonation/deprotonation equilibria that drive charge fluctuations during the simulation. Within Faunus (version 2.16.3), the protonation states of each titratable bead were treated as stochastic variables subject to MC moves that emulate the reversible acid dissociation reaction. Proton concentration is implicitly modelled through the input proton activity, corresponding to the pH, which modulates the acceptance probabilities of protonation changes, in accordance with the pKa values and the local electrostatic environment. By iteratively applying titration moves alongside standard configurational MC steps, the system equilibrates to a charge distribution consistent with the imposed pH and IS. After 10.000 MC sweeps, the protonation states reached a steady-state distribution, allowing for an accurate estimation of the net charge and charge heterogeneity of the mAb. The net charge of mAb1 was then calculated as a function of pH and IS (Figure S8).

A second CG, 174-bead model was developed, by grouping every seven aas of the primary sequence, into a single bead, whose individual diameter (*σ*_*i*_) was determined using equation 1 from the Mwof the seven residues (Figure S14). Each bead was assigned a net charge equal to the average charge of its constituent residues. To assess the validity of this simplified representation, we compared the form factor calculated from the amino acid-level CG model with that of the 174-bead model. As shown in Figure S8, the two models exhibit strong agreement, indicating that the simplified model preserves essential structural features. This reduced model was primarily employed to investigate the collective behavior of mAb1 through large-scale simulations involving multiple mAbs.

### Two-mAb simulations

Using the aa representation of mAb1, we performed two-mAb MC simulations to catch the mAb behavior in the dilute a regime. A first mAb was initially placed in the center of a simulation box, while a second was placed 150 Å away along the **z**-axis. The first mAb could only rotate around its center of mass, while the second could further translate along **z** (Figure S15). mAbs were treated as rigid bodies and interacted with each other through a non-bonded bead-bead potential, *U*_*ij*_ which accounts for both electrostatic and short-range interactions:

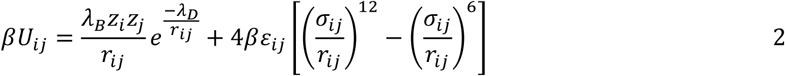

where *λ*_*B*_ = *βe*^2^⁄4*πε*_0_*ε*_*r*_ is the Bjerrum length, with *ε*_0_ being the vacuum permittivity, the relative permittivity of water (being the solvent treated implicitly), *ε*_*r*_ = 80, *e* the electron unit charge, and *β* = 1⁄*k*_*B*_*T* the inverse thermal energy, where *k*_*B*_ is Boltzmann’s constant and the temperature *T* =300 K. *zi* and *zj* are the charges of the *i^th^* and *j^th^* bead, and *r*_*ij*_ the distance between them. The bead-bead minimal approach distance, *σ*_*ij*_, was obtained from equation 1 as *σ*_*ij*_ = (*σ*_*i*_ + *σ*_𝑗_)/2, while the energy depth of the Lennard Jones interactions, *ε*_*ij*_, was fixed to 0.15 kJ/mol to reproduce a mild attraction. The values for *σ*_*ij*_ are reported in Table S1. The Debye length, *λ*_𝐷_, which accounts for salt screening effect (treated implicitly), is calculated through the relation:

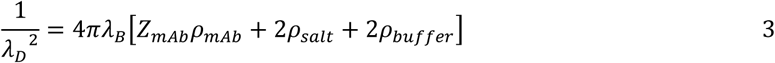

where 𝜌_*mAb*_ is the mAb number density (which is zero for two-mAb simulations as we are assuming infinite dilution), 𝑍_*mAb*_ is the net charge, and 𝜌_*salt*_ and 𝜌_*buffer*_ are, the number densities of salt and of the dissociated buffer ions. Using the simulation setup shown in Figure S15, the radial distribution function of mAb1, g(r), was sampled, from which, the potential of the mean force (PMF), 𝑊(*r*), can be calculated:

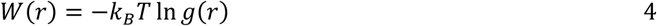

The PMF was used to calculate the reduced second osmotic virial coefficient *B*_2_^∗^, which is defined as:

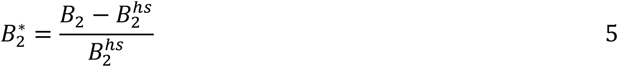

where: 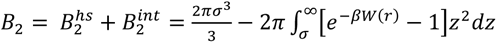 (69). *B*^ℎ𝑠^ represents the hard-sphere contribution to the B2 that arises from steric repulsion after approximating the mAb with a sphere of radius equal to the mAb radius of gyration. *B*_2_^*int*^ is the contribution due to the effective interactions between the mAbs.

With two-mAb simulations, mAb-mAb electrostatic interactions were further characterized. The exact electrostatic energy between two mAbs is the sum of the pairwise Coulomb interactions between all charged beads:

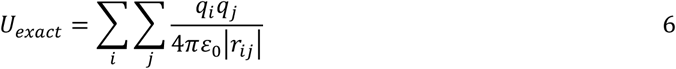

where *r*_*ij*_ is the separation vector between beads *i* and *j,* belonging to the two mAbs. *U*_*exact*_ can be approximated by the sum of multipole moments. More specifically, it can be written as a sum of ion-ion, ion-dipole, dipole-dipole, ion-quadrupole, and higher order terms:

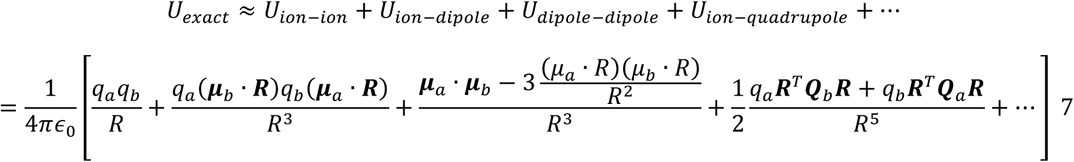

where 𝑞_𝑎_/𝑞_𝑏_, 𝝁_𝑎_/𝝁_𝑏_and 𝑸_𝑏_/𝑸_𝑎_ are net charges, dipole, moments and quadrupole moment tensors of the first and second mAb (a and b) and 𝑹 is the vector connecting their centers of mass.

### Many-mAb simulations

Using the 174-bead model, MC simulations of many mAbs were performed, to reproduce the SAXS scattering intensity of high concentration samples. We used 100 rotating and translating mAbs moving inside a cubic box, whose size was chosen to reproduce a given mAb concentration (Figure S16). In this setup, mAbs still interacted with each other through the bead-bead potential from equation 2.

The intensity at a given scattering vector **q** is then computed according to:

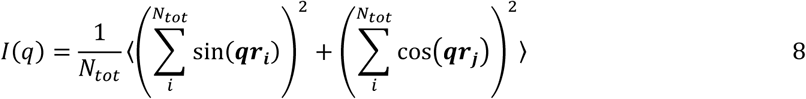

where 𝑁_*tot*_ is the total number of scatterers in the system, defined as the product of the number of mAbs and the number of beads per mAb, and 𝒓_𝒊_ and 𝒓_𝒋_ identify the position of the *i^th^* and *j^th^* bead. The average is performed over a set of scattering vectors, systematically generated by permuting the crystallographic index of the cubic box [100], [110], [111], to define the scattering vector 𝒒 = 2*π*𝑝⁄𝐿(ℎ, *k*, 𝑙), where 𝑝 = 1,2, …, 𝑝_*max*_, and 𝑝_*max*_ = 25.

### Stirring stress protocol

Selected mAb formulations were prepared as 1 mL solutions of 45 mg/mL mAb1 and filtered through a 0.1 μm Whatman Anotop filter (GE Healthcare, Cat no: 6809-2012). The samples were then divided in two 500 µL aliquots and added to separate 4 mL glass screw vials (45X15 mm, MD Scientific, VF013-1545) with a 5x2 mm cylindrical PFTE-coated stirring bar (BOLA, C 350-03). The vials were capped (Thermo Scientific, B7815-13) and sealed before being placed vertically on a magnetic stirring plate (IKA RO 15) and stirred at 1000 rpm overnight (16h). Aliquots of the samples before stirring were diluted 10 times in 150 mM L-arginine HCl in 10 mM histidine pH 6.5 and their concentration measured.

### Monomer recovery after stirring

After stirring, L-arginine HCl in 10 mM histidine pH 6.5 was added directly to the vials at a final concentration of 150 mM. The vials were gently swirled and 400 µL were transferred to microcentrifuge tubes before 2 hours centrifugation at approximately 21000 *g*. Aliquots of the sample supernatants after stirring and centrifugation were diluted 10 times in 150 mM L-arginine HCl in 10 mM histidine pH 6.5 and their concentration measured, before being divided by the initial sample concentration (see previous section) to obtain what we called the monomer recovery here.

## Acknowledgments

The authors acknowledge Novo Nordisk A/S for project funding and providing mAb material. Tobias Palle Holm from the R&ED department at Novo Nordisk A/S is thanked for granting access to the microscope used in this study. The authors also thank Gaetano Invernizzi for mediating the collaboration with the University of Cambridge and general input. Tomas Sneideris is thanked for helping with the analysis of PhaseScan data. Christian Buch Parsbæk is also thanked for his comments to the experiments and research paper. V.F. I.B. S.L. and M.P. acknowledge the support from the VILLUM FONDEN by the Villum Young Investigator Grants “Protein superstructure as Smart Biomaterials (ProSmart)” 2018-2023 (project number: 19175) and “Protein Phase Separation and Solid Transition in Synthetic Cells (ProSeC) 2024-2027 (project number: 53132).

V.F. I.B. and S.L. also acknowledge the Novo Nordisk Foundation (projects NNF20OC0065260 and NNF22OC0080141) for financial support. The authors acknowledge the use of the MAX IV Laboratory for time on the CoSAXS beamline under proposal number 20231488. Research conducted at MAX IV, a Swedish national user facility, was supported by the Swedish Research council under contract 20188-07152, the Swedish Governmental Agency for Innovation Systems under contract 2018-04969 and Formas under contract 2019-02496. The authors further acknowledge DESY research center (Hamburg, Germany), for the provision of experimental facilities and time at PETRA III using beamline P22 (proposals 1188 and 1279).

## Supporting Information

**Figure S1:**
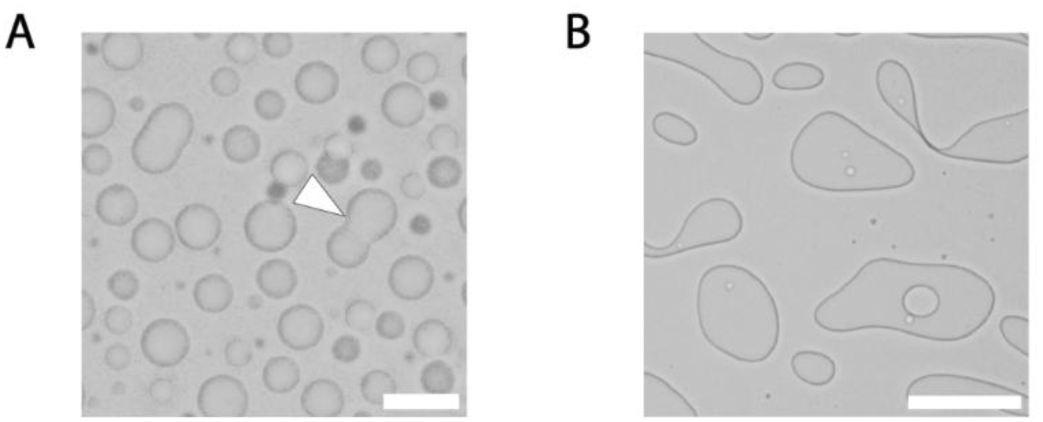
mAb1 LLPS is liquid-like. Brightfield microscopy images of a 45 mg/mL mAb1 sample containing 10 mM NaCl at pH 6.5. The resulting phase-separated droplets are highly dynamic: (A) They fuse and readily coalesce upon contact as indicated by the white arrowhead (10 µm scalebar) and (B) wet the glass surface of the microscopy slide within seconds, emphasizing their liquid-like nature (100 µm scalebar)

**Figure S2:**
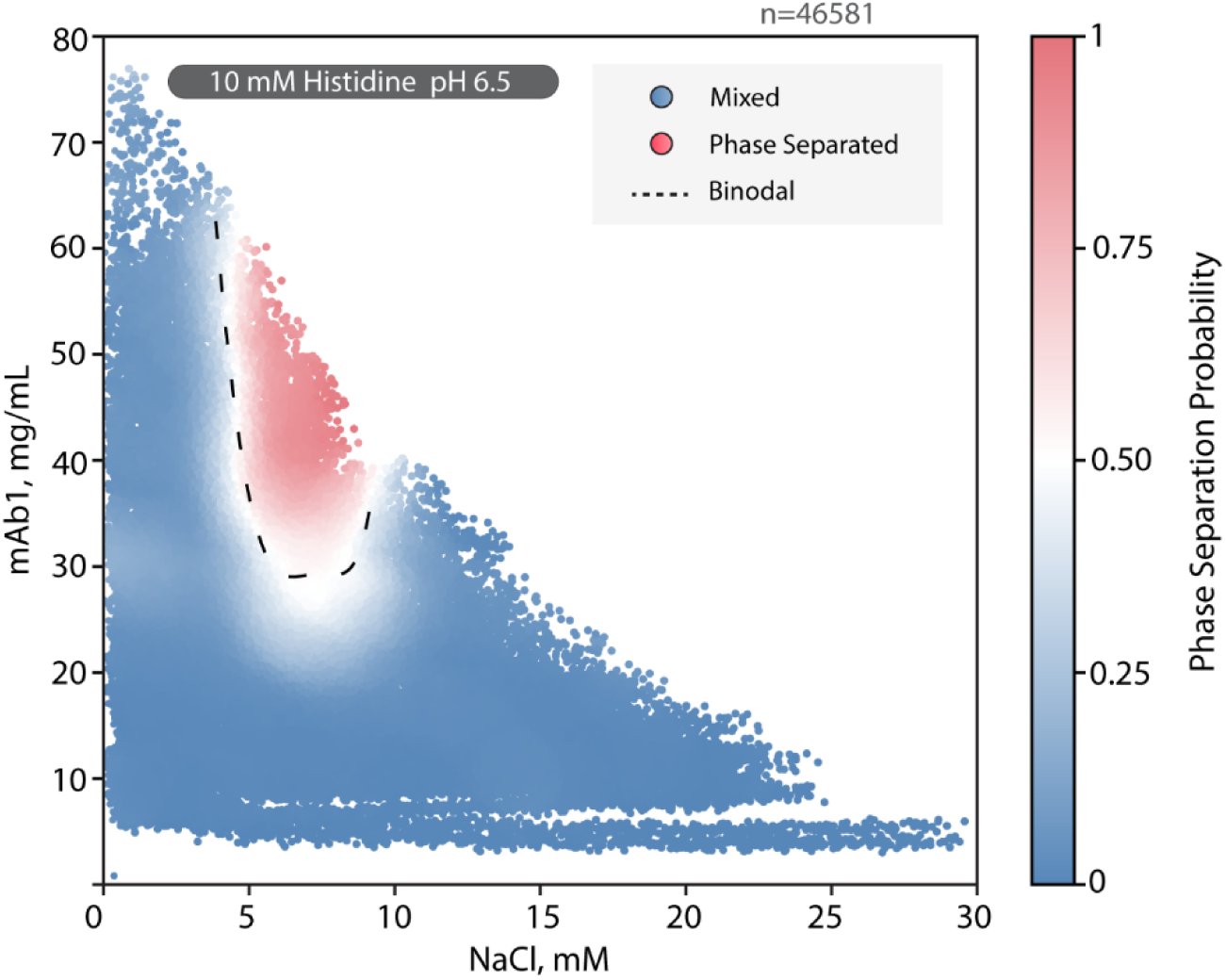
Coexistence curve of mAb1 as a function of NaCl determined using combinatorial microfluidics. Each of the 46581 observations is shaded according to phase separation probability based on local radial averaging. An eye guide for the approximate location of the binodal is indicated by the dashed black line.

**Figure S3:**
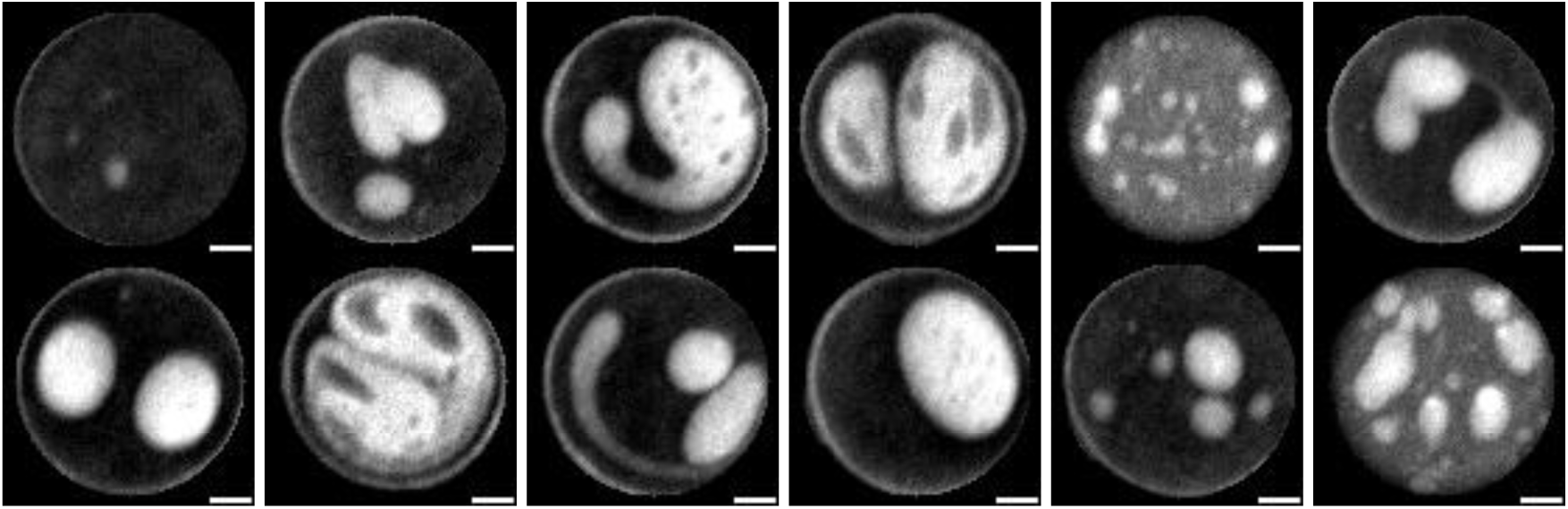
Selected dense phase morphologies detected with PhaseScan using fluorescence microscopy (20 µm scalebar)

**Figure S4:**
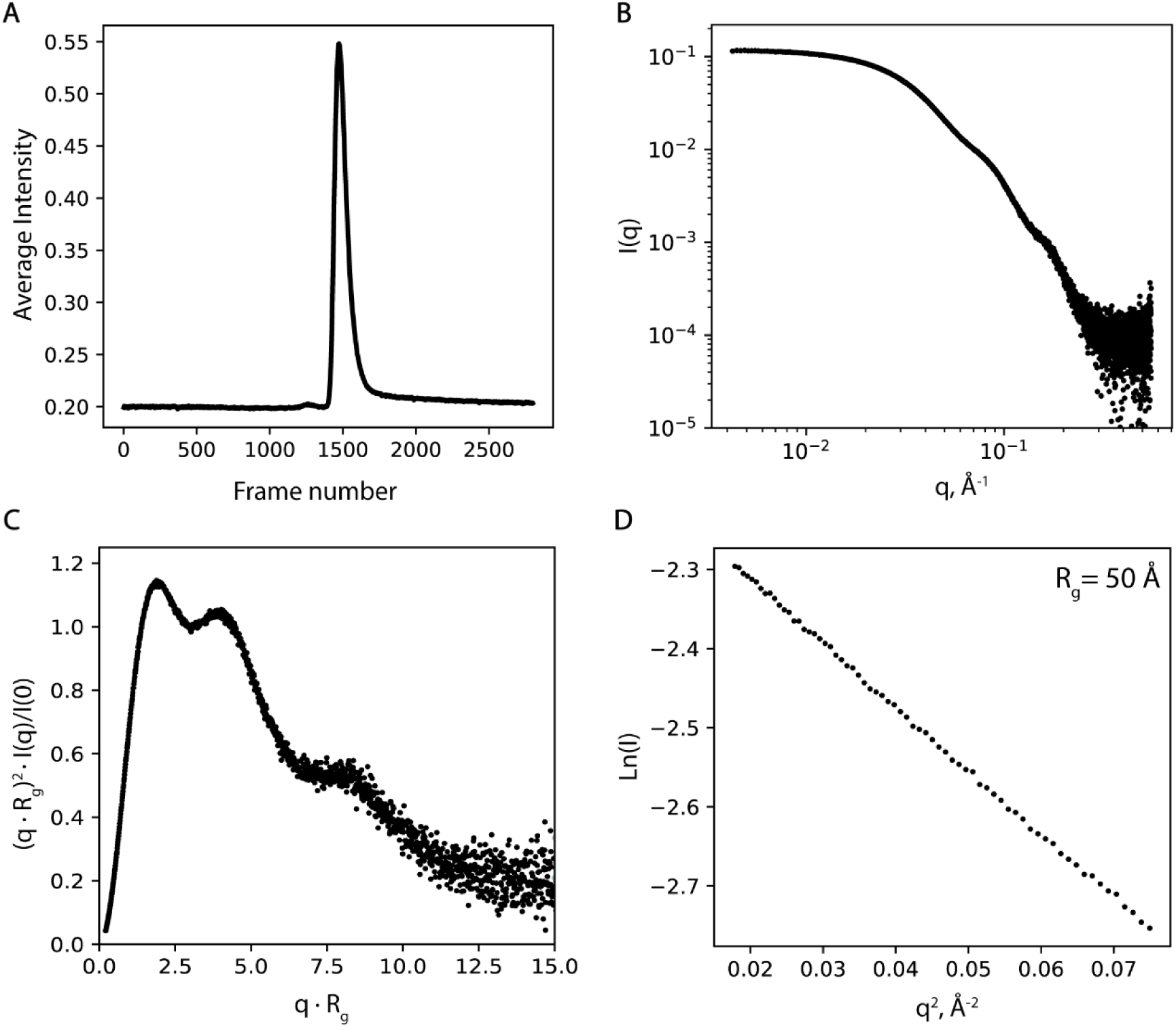
Overview of SEC SAXS measurements of mAb1. (A) Elution profile of 10 mg/mL mAb1 in 10 mM histidine pH 6.5 + 150 mM NaCl mobile phase showing the average integrated SAXS intensity, I(q), (1.48*10^-3^ Å^-1^ < **q** < 0.556 Å^-1^) as a function of frame number (B) Average scattering curve of frames 1454 to 1521, corresponding to monomeric mAb1 (highest peak of the elution profile in A) (C) Dimensionless Kratky plot showing three peaks, as expected for IgG molecules (D) Guinier plot for the monodisperse mAb1 sample showing linearity in the Guinier range and the radius of gyration determined from the slope.

**Figure S5:**
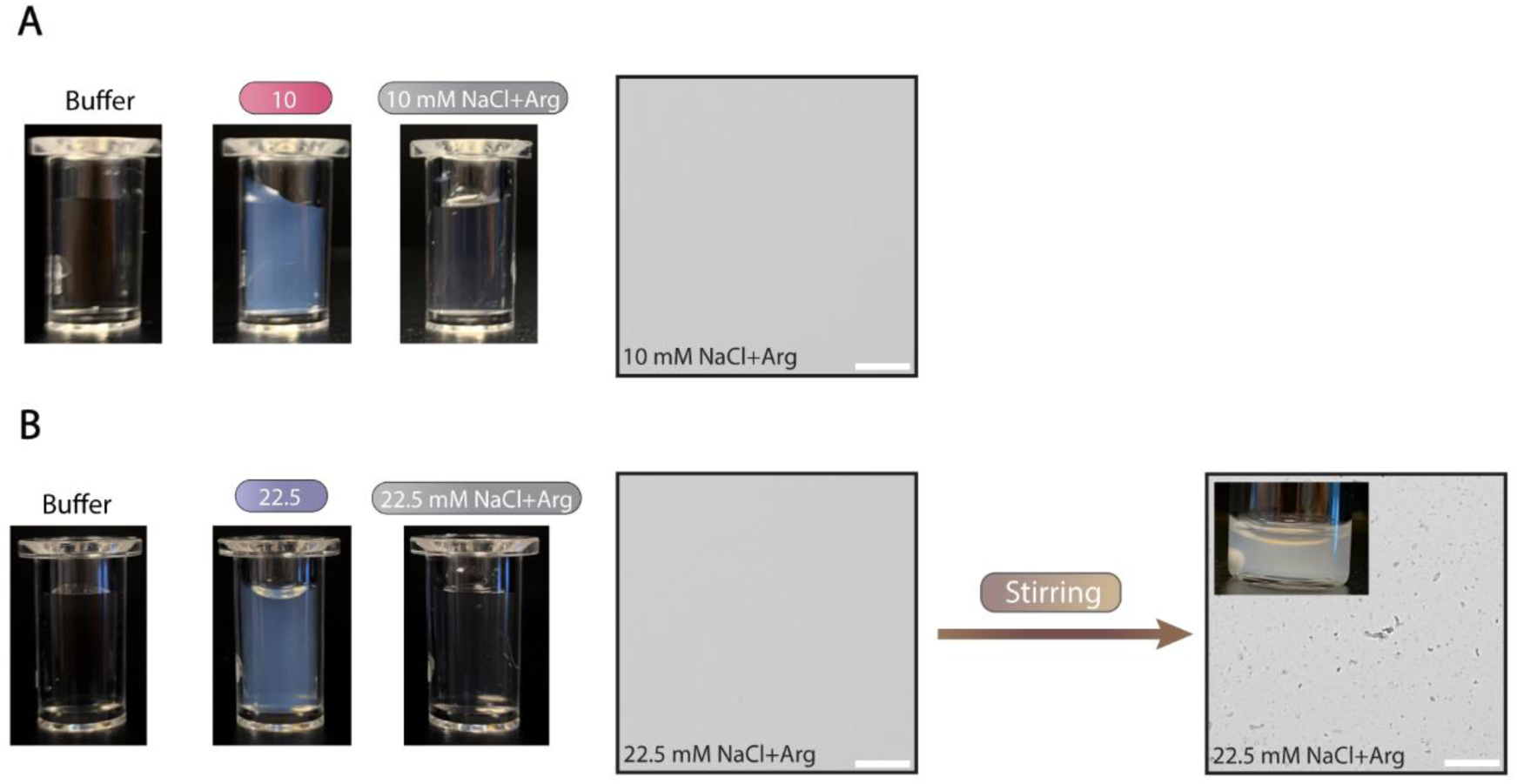
Suppression of LLPS and opalescence with L-arginine HCl and effect of stirring. (A) Left: Visual appearance of an LLPS sample containing 45 mg/mL mAb1 in 10 mM histidine, 10 mM NaCl, pH 6.5 compared to 45 mg/mL mAb1 in 10 mM histidine, 10 mM NaCl, 150 mM L-arginine-HCl, pH 6.5 (10 mM NaCl+Arg). The latter was added prior to adding the NaCl that would have otherwise made the sample undergo LLPS. Right: Brightfield microscopy of the 10 mM NaCl+Arg sample showing that no LLPS is detected under these conditions. The scalebar represents 50 µm. (B) Same setup as in (A) but with the 22.5 mM (opalescent sample) as well as images of the effect of stirring at 1000 rpm (16 hours), showing the increased visual turbidity of the sample and the presence of aggregates in the microscopy image. The scalebars represents 50 µm.

**Figure S6:**
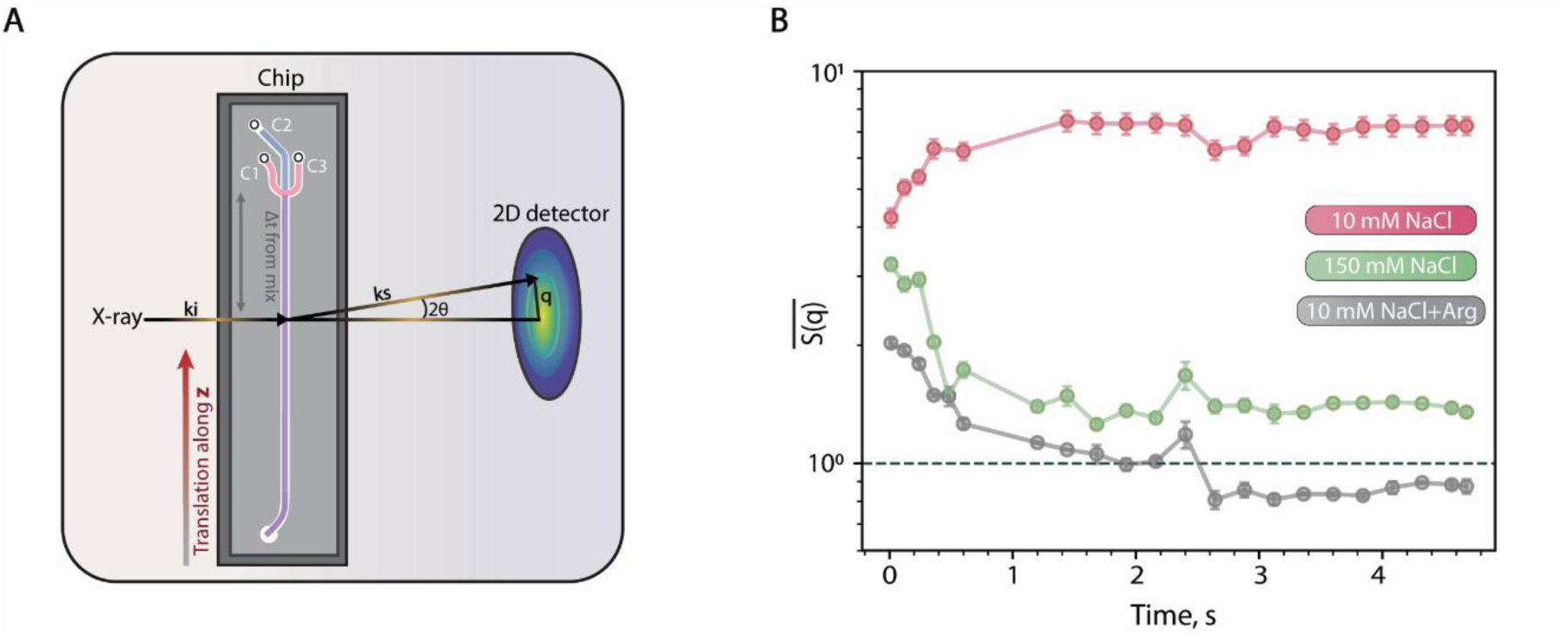
(A) Schematic representation of the microfluidic chip coupled to the Co-SAXS beamline recorded at 22 distances from the mixing point across the channel. The distances were converted to timepoints here. Data was recorded at a total flowrate of 60 µL/min. (B) Averages of the first three S(**q**) values, that is, the first three datapoints at low **q** of each of the 22 curves from the time-resolved experiment, converted to timepoints here.

**Figure S7:**
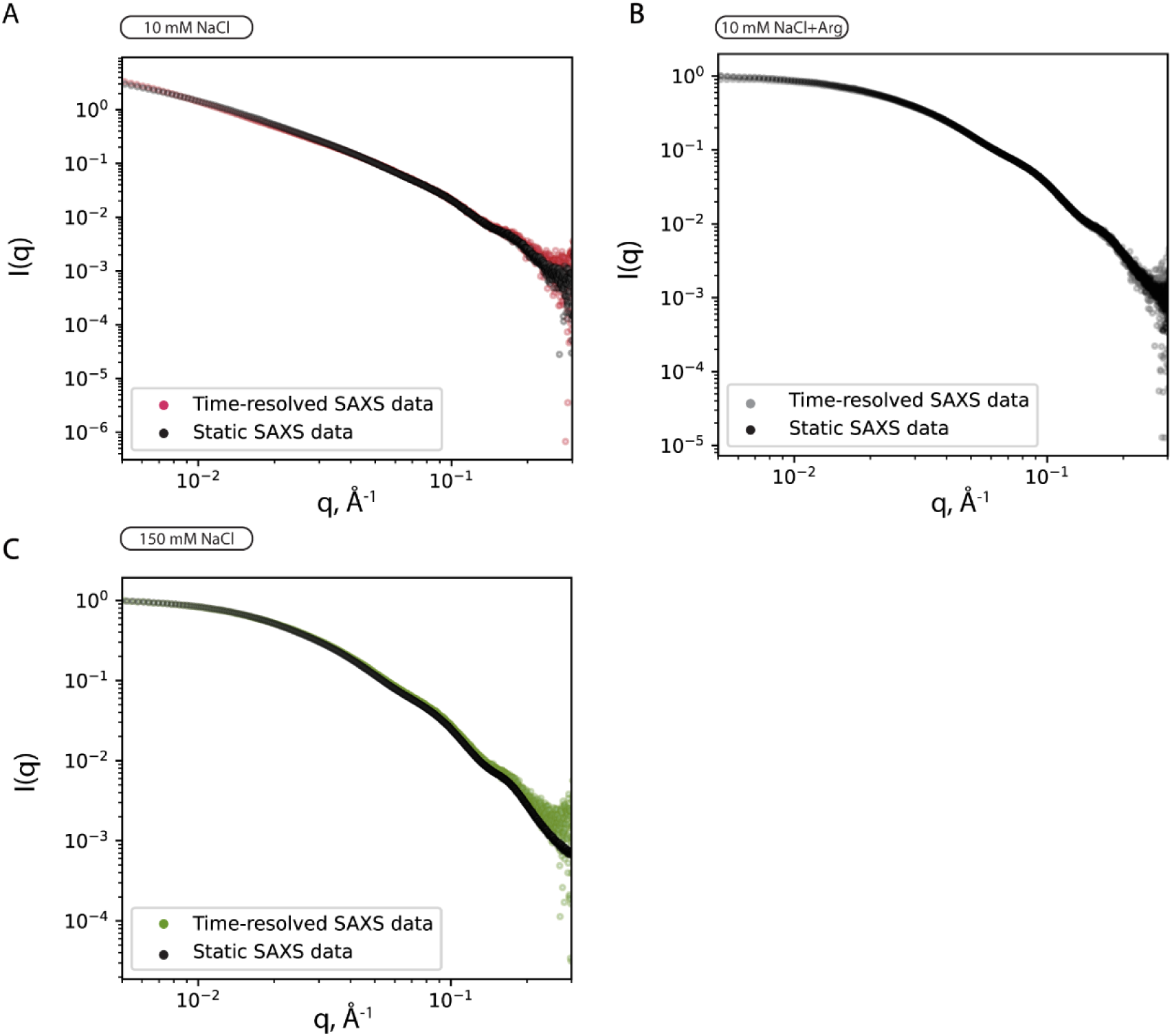
Overlay of the scattering curves (intensity I(**q**) vs **q**) for the last timepoint in the time resolved SAXS experiments (colored curves) and scattering curves from the static SAXS experiment (black curves) for 45 mg/mL mAb1 in 10 mM histidine pH 6.5 containing (A) 10 mM NaCl, (B) 10 mM NaCl + 150 mM arginine HCl (10 mM NaCl+ Arg) or (C) 150 mM NaCl.

**Figure S8:**
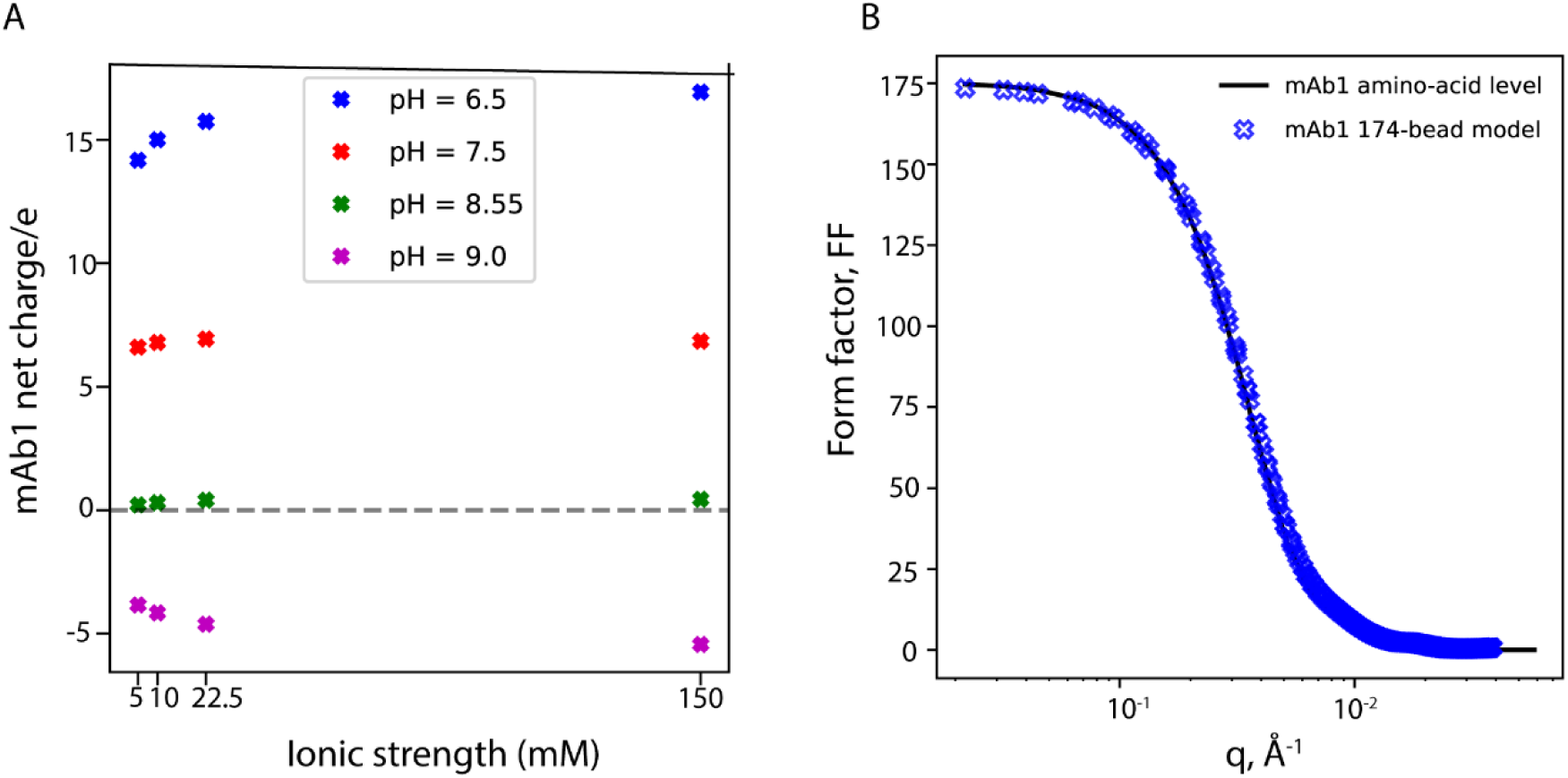
(A) mAb1 net charge in electron units at pH 6.5 (blue), pH 7.5 (red), pH 8.55 (green), and pH 9.0 (purple) as a function of solution IS. (B) mAb1 form factor comparison between the structure coarse-grained at the amino acid level (blue full line) and the 174-bead model (blue symbols).

**Figure S9:**
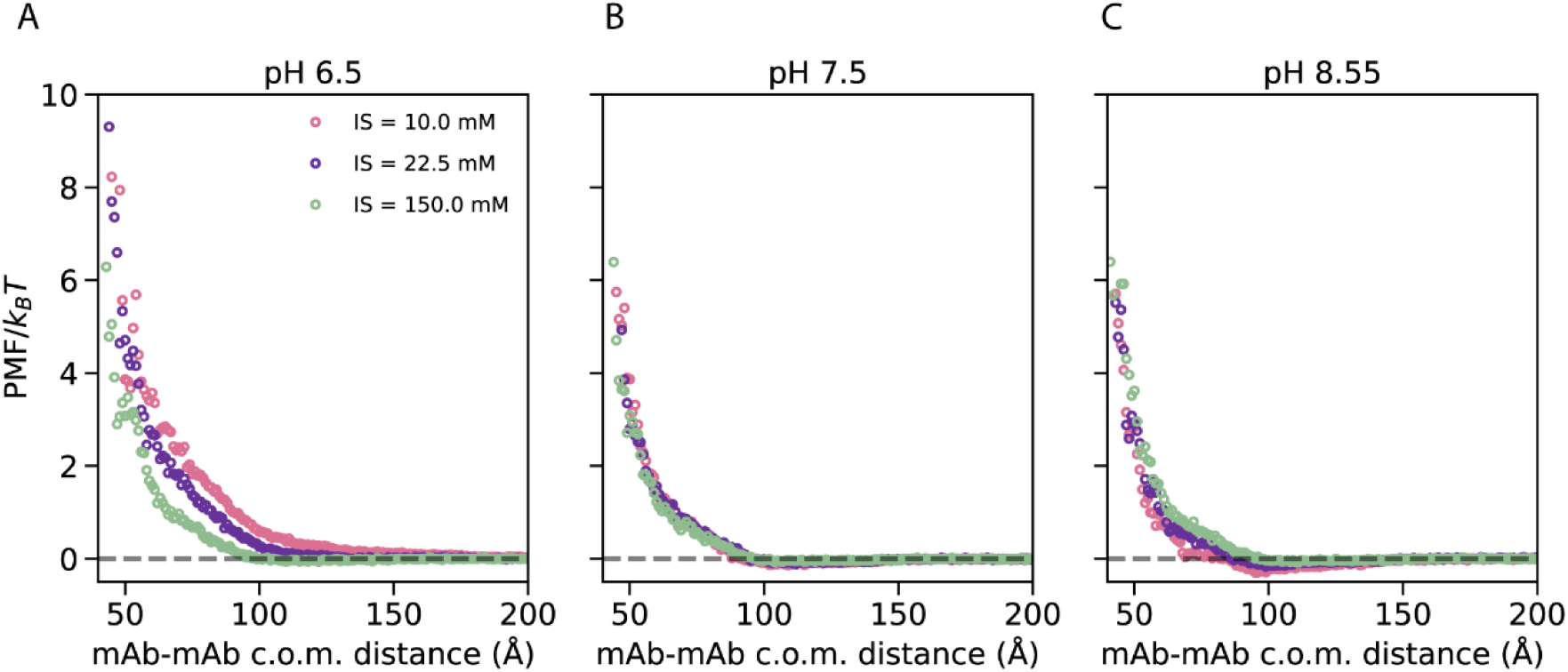
PMF analysis. MAb1 PMF for pH (A) 6.5, (B) pH 7.5, and (C) pH 8.55, for ionic strength conditions of 10 mM (pink), 22.5 mM (purple), and 150 mM (green). Here, *ε*_*ij*_ was set to 0.15 kJ/mol to simulate a mild attraction between mAbs.

**Figure S10:**
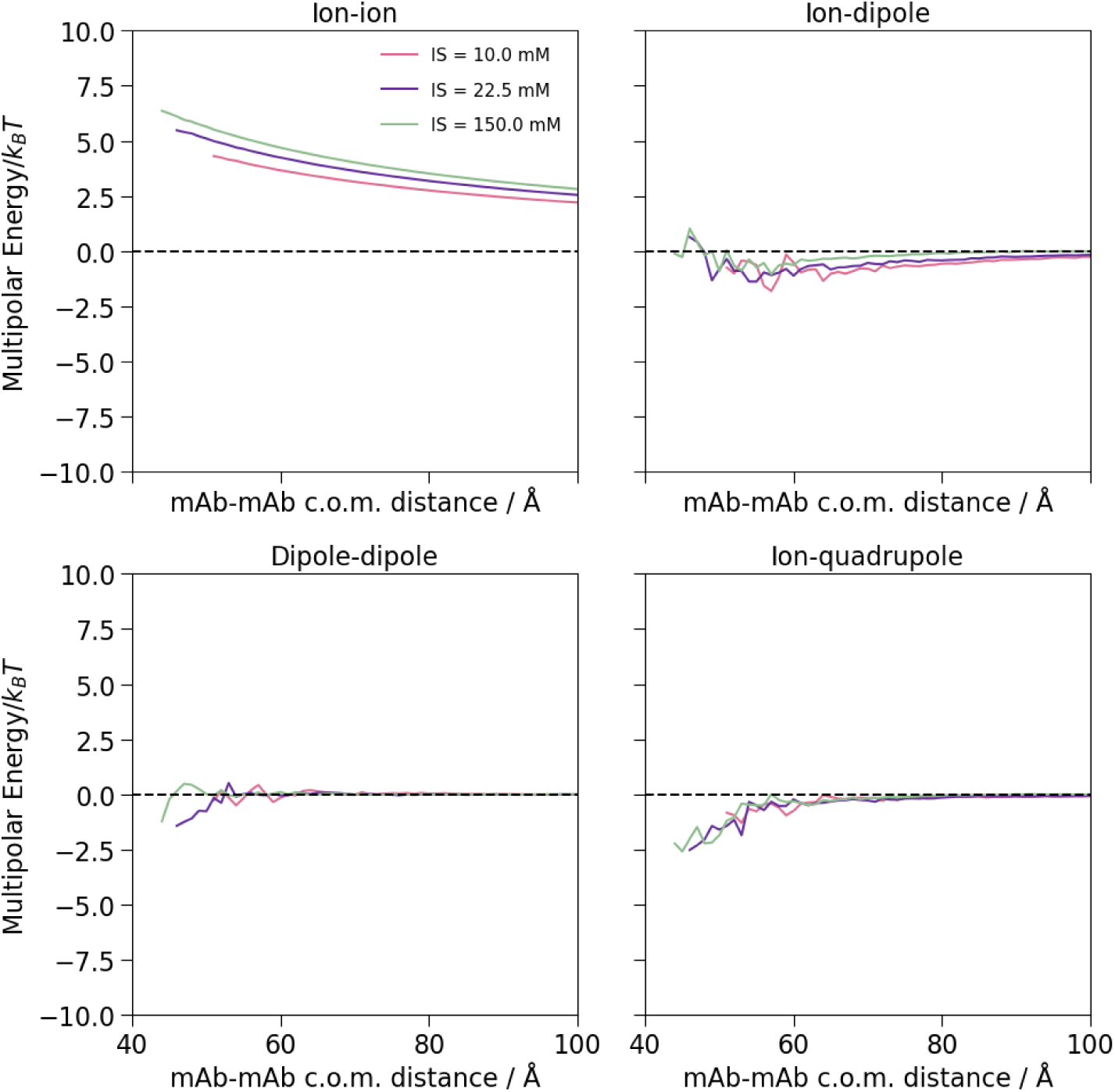
mAb1 multipole expansion contributions for pH = 6.5 resulting from two-mAb simulation where *ε*_*ij*_ was set to 0.15 kJ/mol to simulate a mild attraction between mAbs.

**Figure S11:**
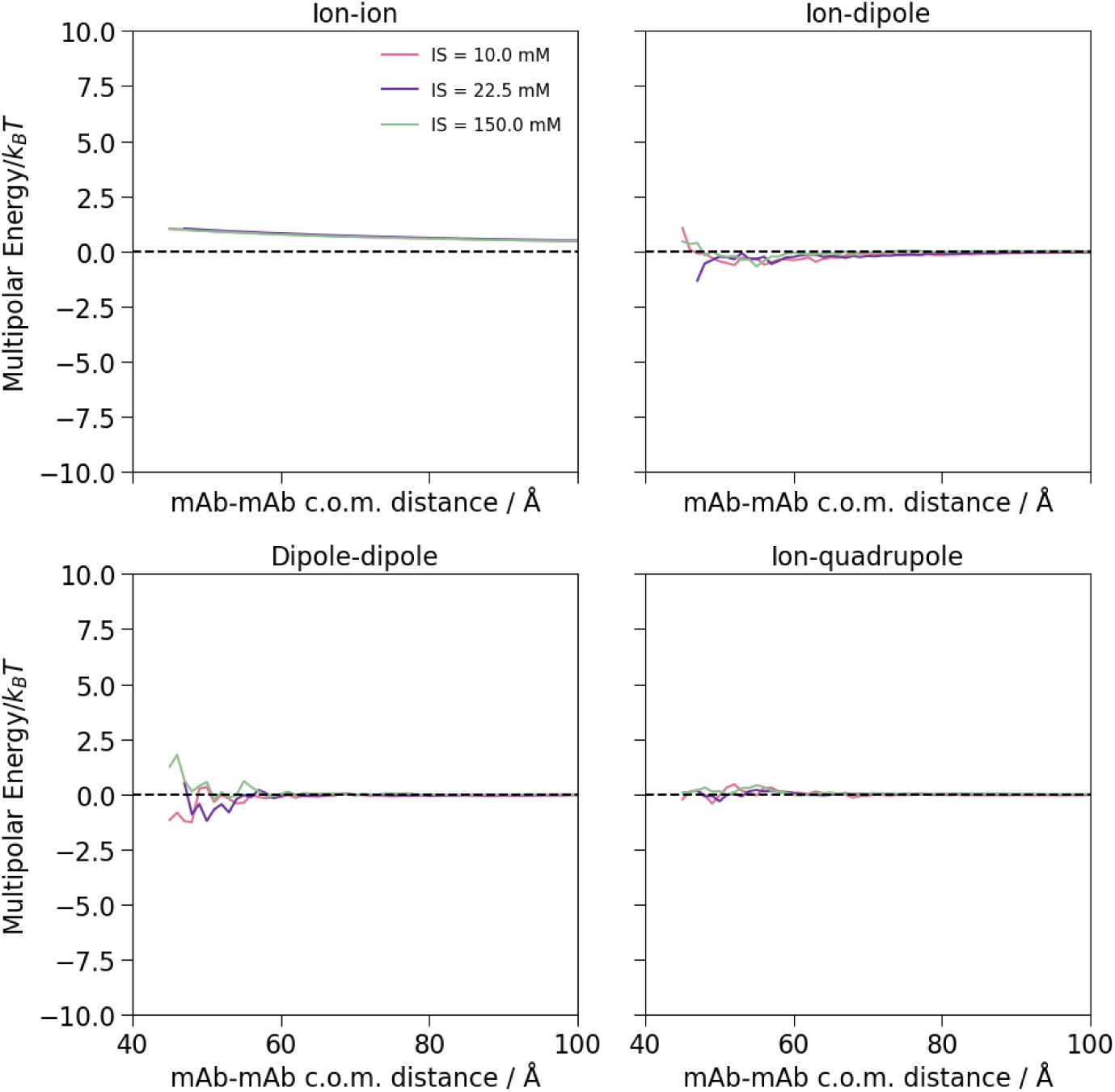
mAb1 multipole expansion contributions for pH = 7.5 resulting from two-mAb simulation where *ε*_*ij*_ was set to 0.15 kJ/mol to simulate a mild attraction between mAbs.

**Figure S12:**
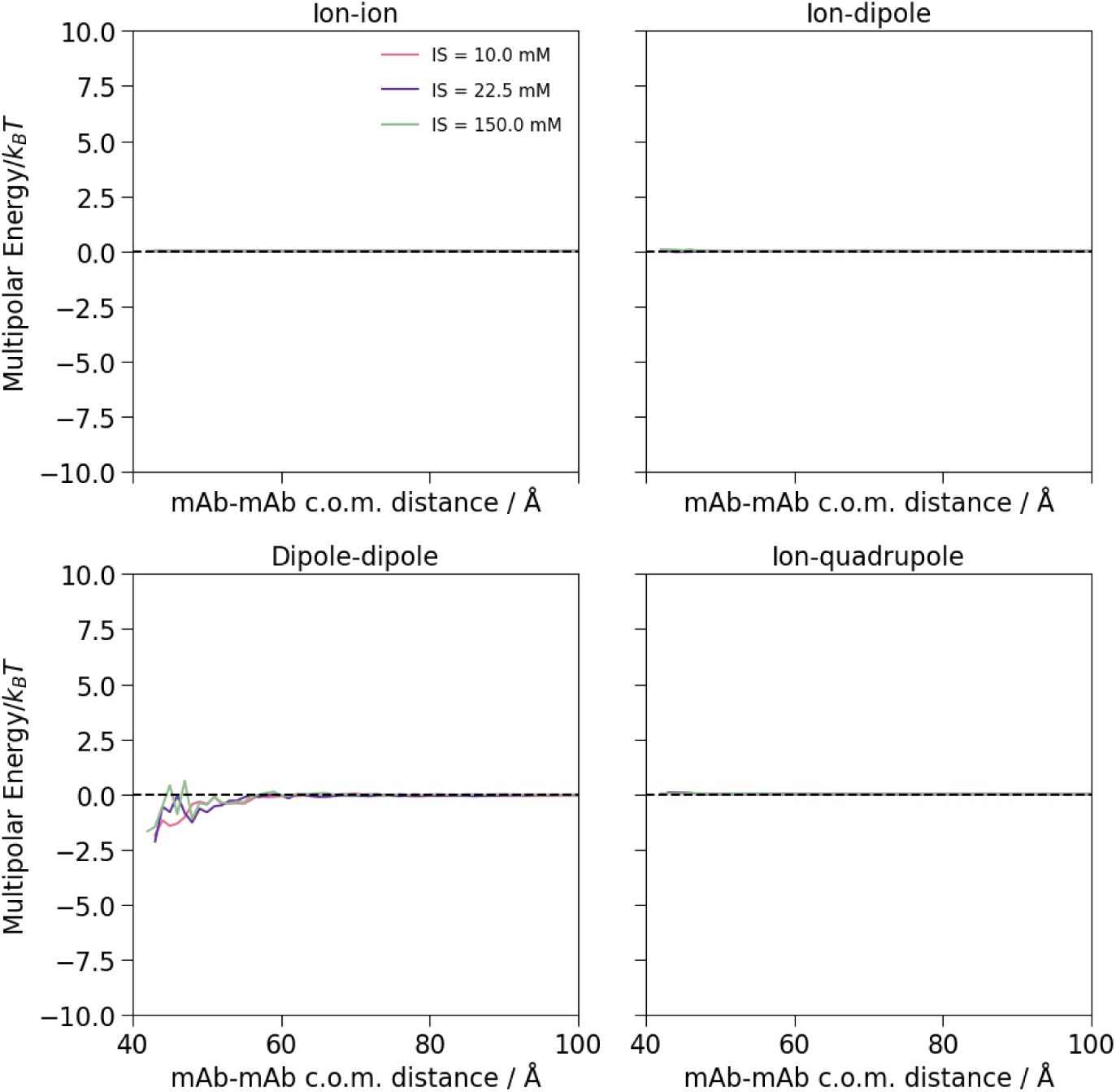
mAb1 multipole expansion contributions for pH = pI = 8.55 resulting from two mAb simulation where *ɛij* was set to 0.1 kJ/mol to simulate a mild attraction between mAbs.

**Figure S13:**
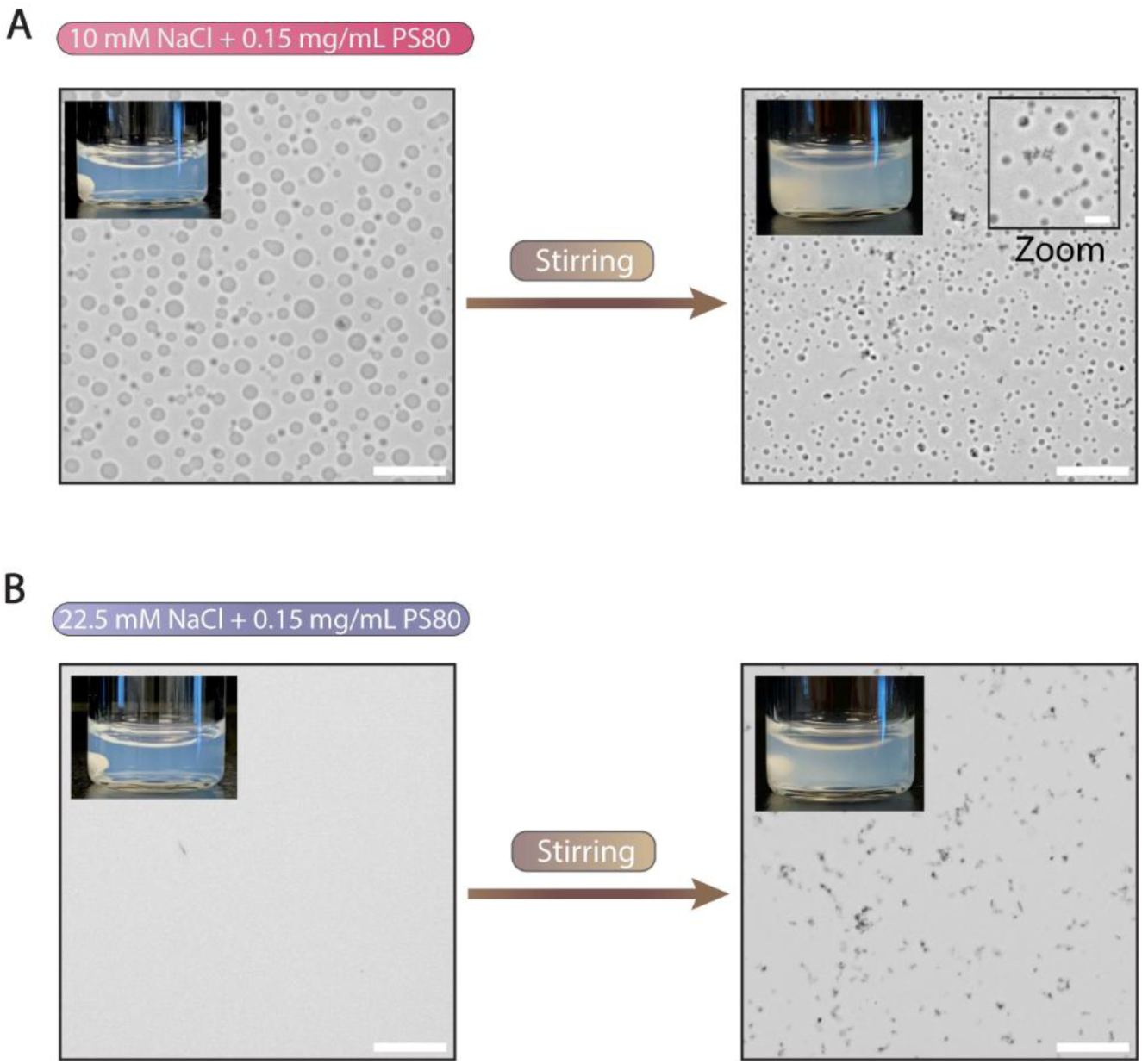
Effect of PS80 on the stirring induced aggregation. (A) Visual appearance and brightfield microscopy images of 45 mg/mL mAb1 in 10 mM histidine pH 6.5 containing 10 (A) or 22.5 (B) mM NaCl with 0.15 mg/mL PS80 added. The left images are before stirring and the images on the right are after overnight stirring as described in the methods section. The scalebar represents 50 µm in the main microscopy image and 10 µm in the inset zoom image. In the 10 mM NaCl + 0.15 mg/mL PS80 images, the difference in droplet size arises from their fast dynamics and difference in times at which the images were acquired.

**Figure S14:**
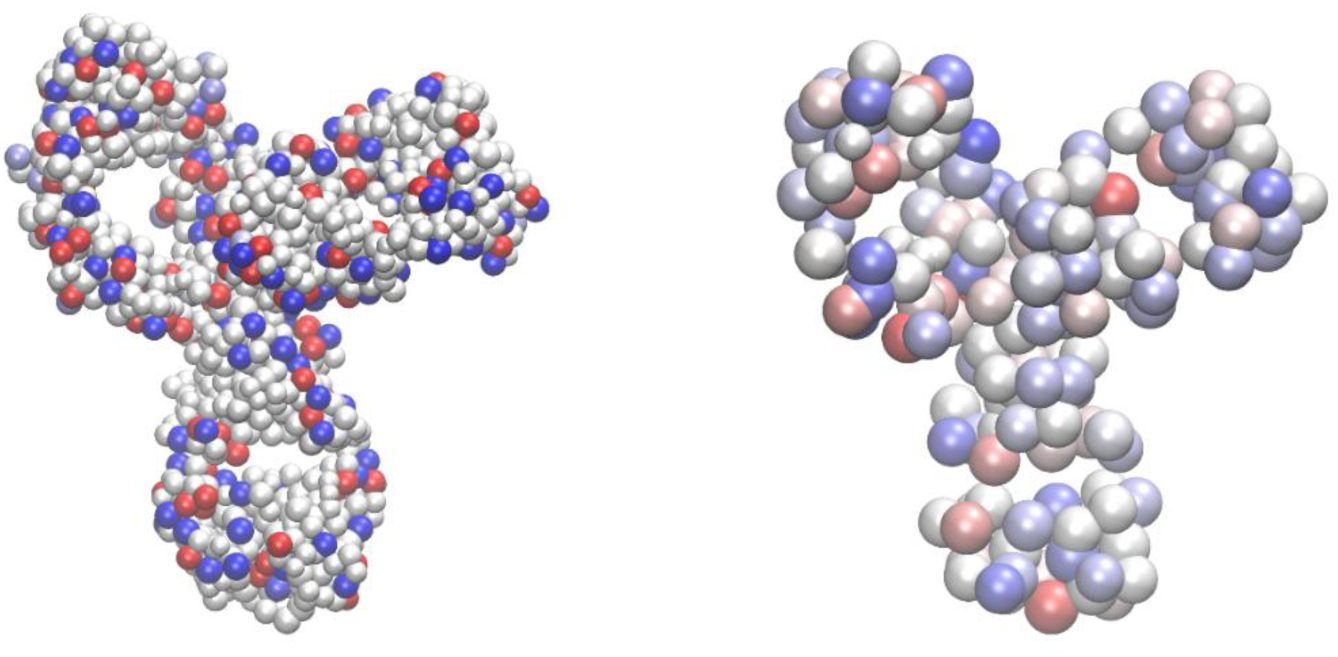
mAb1 coarse-grained models. Amino-acid level (left), and 174-bead model (right). Blue, red, and grey beads represent, respectively, positively, negatively, and neutral charged beads. The current representation reflects the protonation state of the mAb in a solution with 10 mM of NaCl and pH 6.5.

**Figure S15:**
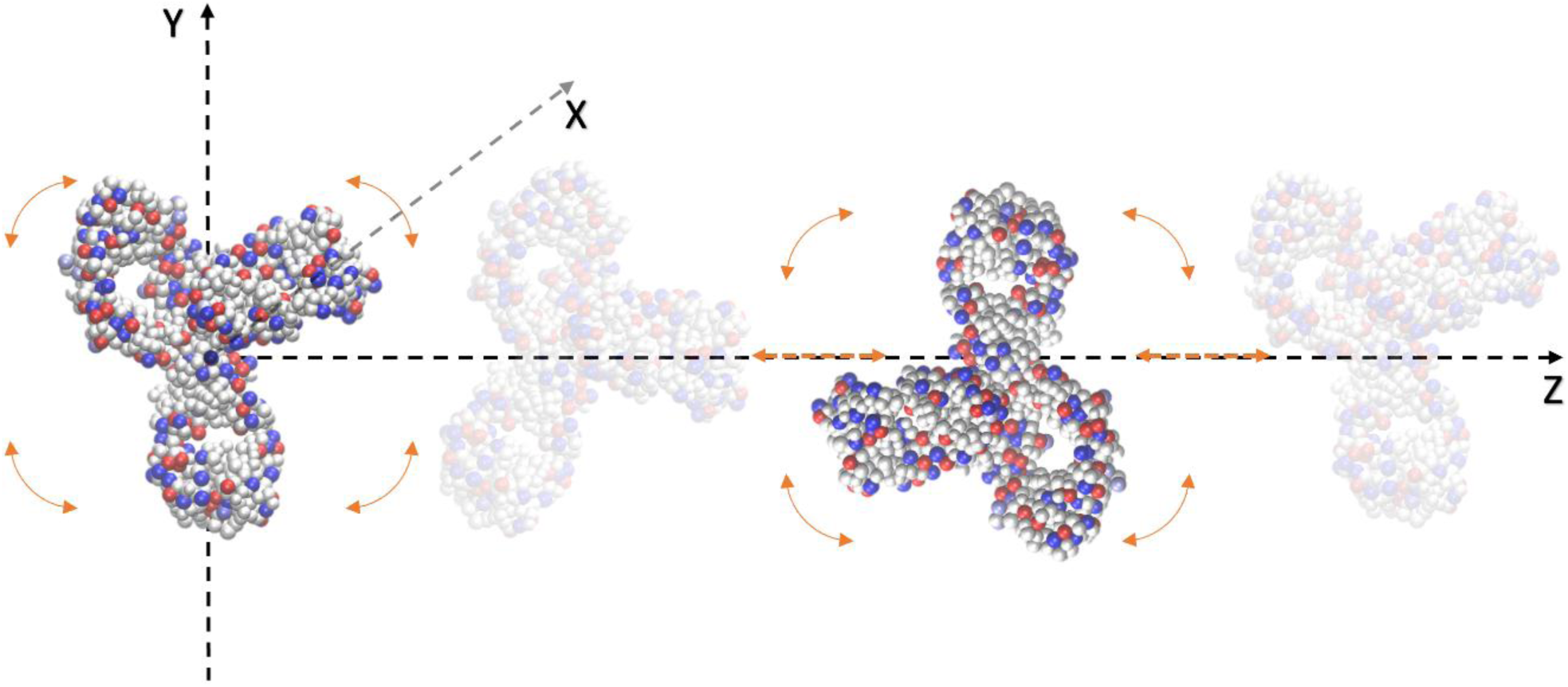
Two-mAb simulations setup. A first mAb (on the left) is placed in the center of the simulation box and can only rotate around its center of mass. A second mAb (on the right) can rotate around its center of mass and rigidly translate along the Z-axis. By sampling different relative positions of the mAbs’ center of masses on the z-axis, we sample the radial distribution function, g(r), and consequently, from equations 4 and 5, the PMF and the reduced second osmotic virial coefficient, *B*^∗^.

**Figure S16:**
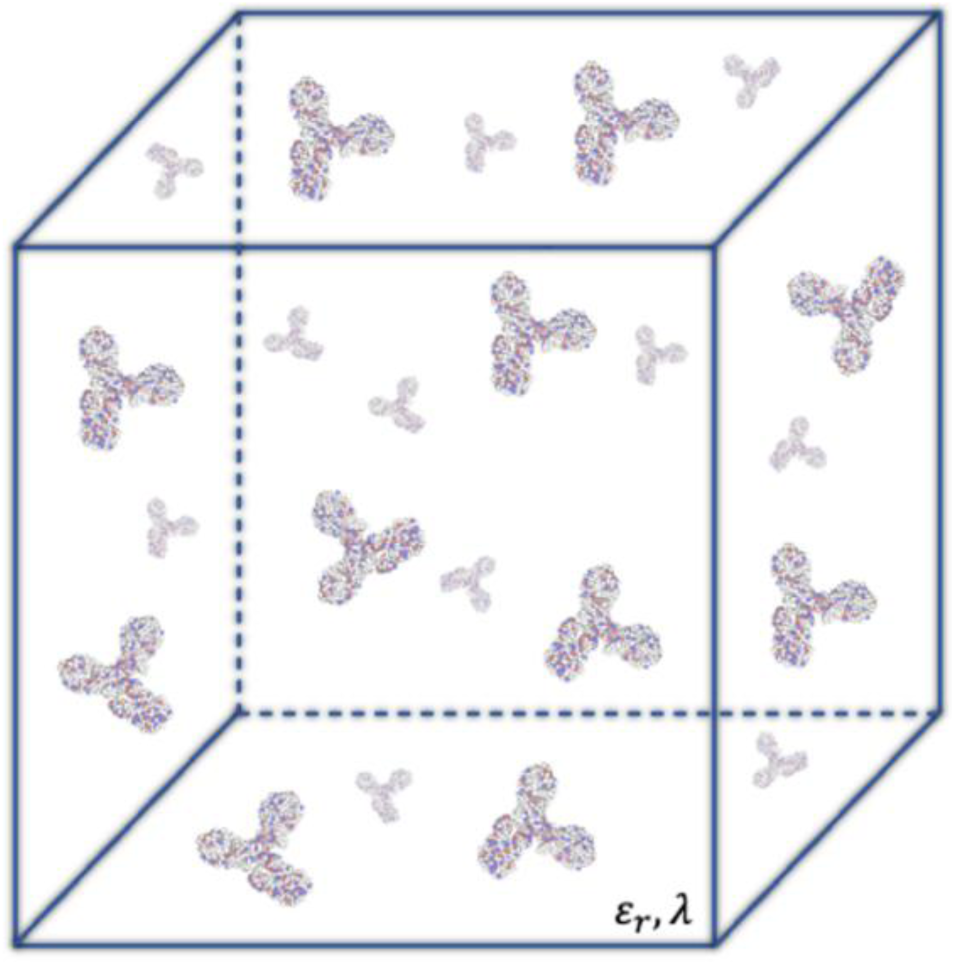
Schematic representation of many-mAb simulations. Inside the simulation box, mAbs can rigidly rotate and translate in all directions. Solvent and salt screening effect are treated implicitly in the interaction potential through the dielectric water constant *ε*_*r*_, and the Debye length, *λ*, respectively (see equation 2).

**TABLE S1:** Bead name and corresponding sigma values for the mAb1 coarse-graining at the amino acid level.

| Bead | Sigma | Bead | Sigma |
| --- | --- | --- | --- |
| CTR | 3.13 | TRP | 6.95 |
| HCTR | 3.13 | GLY | 4.69 |
| NTR | 2.99 | ARG | 6.50 |
| HNTR | 2.99 | HARG | 6.50 |
| ALA | 5.02 | LYS | 6.05 |
| ILE | 5.80 | HLYS | 6.05 |
| LEU | 5.80 | ASP | 5.94 |
| MET | 6.16 | HASP | 5.94 |
| PHE | 6.41 | LASP | 5.94 |
| VAL | 5.56 | GLU | 6.16 |
| PRO | 5.56 | HGLU | 6.16 |
| SER | 5.39 | LGLU | 6.16 |
| THR | 5.64 | GLN | 6.12 |
| TYR | 6.65 | ASN | 5.91 |
| HTYR | 6.65 | HIS | 6.29 |
| CYS | 5.72 | HHIS | 6.29 |
| HCYS | 5.72 | GLC | 6.62 |
| CYX | 5.72 |  |  |

## Notes

### Competing Interest Statement

Competing Interest Statement: N.L, M.N, J.S.M, A.H. and M.G.J are employees of Novo Nordisk A/S.

### Summary of Updates

A molecular simulation study is now included, leading to a revision of the text in all its parts and changes to Figures 3 and 5, and the addition of Figures S8-S12, S14-S16 and Table S1 in the supplementary materials. Authors list is updated.

